# Rapid Podocyte ablation Causes Acute Renal Tubule Cell Necrosis and Interstitial Fibrosis

**DOI:** 10.64898/2026.03.19.712955

**Authors:** Yilin Chen, Mohammad Islamuddin, Xiaofeng Ding, Jefferson Evangelista, Avi Salomon, Gabriella M Hidalgo, Shumei Liu, Cecily C Midkiff, Ryousuke Satou, Jia L. Zhuo, Jay K Kolls, Vecihi Batuman, Rhea Bhargava, Robert V Blair, Xuebin Qin

**Author notes:** Address for correspondence: Xuebin Qin, PhD, Division of Comparative Pathology, Tulane National Biomedical Research Center, Health Sciences Campus, 18703 Three Rivers Road, Covington, LA 70433, USA. These authors contributed equally: Yilin Chen and Mohammad Islamuddin.

## Abstract

It remains unclear whether podocyte loss directly causes acute renal tubular cell (RTC) damage and interstitial fibrosis, thereby leading to renal failure. Here, we applied intermedilysin (ILY)-mediated human CD59 (hCD59) cell ablation to generate an acute, specific podocyte-ablation mouse model. Cre-induced hCD59 transgenics (ihCD59) were crossed with *Nphs2Cre* to generate *ihCD59^+/-^/Nphs2Cre^+/-^* mice. The specific and rapid podocyte-ablation mediated by ILY injection directly caused RTC necrosis, leading to renal failure and even death within 2-3 days in a dose-dependent manner. Treating mice that received an ILY lethal dose with peritoneal dialysis or administering a non-lethal dose, we extended their survival beyond six weeks and found that mice developed interstitial fibrosis and glomerulosclerosis with persistent proteinuria and tubule damage. Podocyte-ablation caused massive disruption of glomerular function at week 1, and then partial recovery by week 2. Genes and pathways of TLRs and apoptosis, and mitochondrial functions were respectively upregulated and downregulated in both ablated-podocyte mouse and biopsied-glomerulonephritis patient kidney samples. Together, this rapid podocyte-ablation causes acute RTC necrosis that progresses to interstitial fibrosis in this mouse model, which is applicable for dissecting mechanisms underlying podocyte injury-mediated tubular damage and glomerular repair, with the potential to reveal novel therapeutic targets for kidney diseases.

## Introduction

Podocytes (Pod) play a vital role in kidney health as damage to Pod results in significant proteinuria, often occurring with associated damage to renal tubule epithelial cells (RTC). The relationship between Pod and RTC injury is a complex, incompletely understood aspect of kidney diseases. Pod loss and associated RTC damage and/or interstitial fibrosis are implicated in the pathogenesis of nephrotic syndrome (NS), autoimmune related nephropathy (ARN), and diabetic nephropathy (DN). However, whether Pod injury or loss directly causes acute RTC damage and interstitial fibrosis, thereby leading to renal failure remains unclear and has not been investigated extensively. The mechanisms underlying Pod damage-mediated acute RTC necrosis and interstitial fibrosis remain largely unknown. Primarily due to the lack of established experimental tools capable of dissecting the effect of Pod damage from RTC damage and interstitial fibrosis. Addressing these questions is critical for us to better understand Pod biology and develop efficient targeted treatments for kidney diseases.

Previously, diphtheria toxin (DT)/diphtheria toxin receptor (DTR) method has been used to conditionally damage Pod cells in mice and rats. Those studies have demonstrated that DT-mediated injury causes a dose-dependent loss of Pods, which can lead to glomerulosclerosis and progression to end-stage kidney disease (ESKD)(1–3). Ablation of developing Pods through transgenic expression of DT on Pod disrupts cellular interactions and nephrogenesis both inside and outside the glomerulus(4). It has also been reported that treatment of wild-type C57BL/6 mice with DT leads to marked transient, and completely reversible proteinuria, as a consequence of Pod dysfunction that is morphologically characterized by foot process fusion and detachment from the glomerular basement membrane(5), which indicates that DT alone has toxic effect on Pods in WT mice. In addition, significant kidney damage was demonstrated in another mouse model (called POD-ATTAC mice) using the human podocin promoter (Nphs2) to drive expression of a mutant FKBP fusion protein that, once dimerized, induces physiologic, caspase-8–mediated apoptosis in Pods. The kidney damage in this model mimicked several aspects of human renal disease, such as foot process effacement, mesangial expansion, and glomerulosclerosis (6). These models, along with many other genetic mutation models for nephrotic syndrome, models for ARN, and other toxin-induced models of Pod damage, have provided essential tools for us to better understand Pod biology and test the effectiveness of potential treatments(1, 2, 5–13). However, these tools have not been used to investigate whether Pod damage directly causes RTC injury and interstitial fibrosis, or to elucidate the underlying cellular mechanisms. This may be due to these cell ablation models having off-target effects, a narrow pharmacological window, and/or a slow mechanism of action (LPS, chemicals, murine knockout models or diphtheria toxin [DT]/DT receptor [DTR])(2, 3, 14),

To overcome these limitations, we have applied a rapid and specific cell ablation tool, namely, intermedilysin (ILY)-mediated human CD59 (hCD59) (ILY-hCD59) cell ablation in the mice(14–18). To generate a novel, rapid and specific Pod ablation mouse model, Cre induced-hCD59 transgenic line (*ihCD59*) previously (14, 16–18) were crossed with podocin-Cre mice (or *Nphs2Cre*)(19) to generate the *ihCD59^+/-^/Nphs2Cre^+/-^* mice. ILY, a bacterial toxin, exclusively binds to hCD59 but not CD59 from any other species and selectively lyses cells expressing hCD59 in a few seconds(14–18). The ILY injection causes rapid and specific Pod destruction leading to proteinuria, renal failure, and death in the *ihCD59^+/-^/Nphs2Cre^+/-^* but not the *ihCD59^+/-^/Nphs2Cre^-/-^*control mice. The specific and acute Pod damage, degree of RTC damage, and the occurrence of the clinical manifestations are dependent on the dose of ILY. Electronic microscopic analyses demonstrated the rapid loss of Pods and foot processes, which resulted in massive proteinuria, loss of RTCs brush borders, and acute tubular necrosis. At a molecular level, bulk RNA analysis of the kidneys after ILY injection to mice revealed that toll like receptor (Tlr) genes, Tlrs signaling pathways, and necrotic and apoptotic pathways are dramatically increased at both 4 and 7 hours after the Pod ablation. Further, we conducted peritoneal dialysis (PD) in hCD59-expressed Pod mice treated with ILY. We found that PD extended the survival of the mice beyond 6 weeks after ILY injection. Despite this, these mice developed ongoing RTC degeneration and interstitial fibrosis. Together, our results demonstrate that rapid Pod ablation causes acute renal tubule cell necrosis and interstitial fibrosis.

## Results

### Specific Pod ablation causes acute kidney injury (AKI) characterized by hematuria, albuminuria, progressive anuria, and renal failure

*ihCD59^+/-^/Nphs2Cre^+/-^* mice (Experimental mice (Exp)) were generated by crossing Cre induced-hCD59 transgenic line (*ihCD59*) established previously(14, 16–18) with podocin-Cre mice (or *Nphs2Cre*)(19). Immunohistochemical (IHC) analysis for hCD59 demonstrated strong and selective expression restricted to glomerular Pods in *ihCD59^+/-^ /Nphs2Cre^-/-^* mice (Control (Ctrl)) (**Supplementary Figure (SFig) 1a**). In contrast, no hCD59 immunoreactivity was detected in glomeruli of B6 WT or *ihCD59^+/-^/Nphs2Cre^-/-^*control mice (**SFig. 1a**). Of note, a non-specific staining pattern for hCD59 was present in the brush border of some RTCs (black arrows, **SFig. 1a**). The non-specific nature of this staining pattern is supported by its presence in both the B6 WT and *ihCD59^+/-^/Nphs2Cre^-/-^* mice. The specific Cre recombinase induced hCD59 expression on Pods (red arrows, **SFig. 1a)** but not on other cells in the kidney and in other organs (such as liver, brain, heart and gut and spleen, et al) (**SFig. 1b**). Further, immunofluorescence (IF) staining confirmed co-localization of hCD59 with the Pod marker synaptopodin (SYNPO) in glomeruli of *ihCD59^+/-^ /Nphs2Cre^+/-^* mice (**SFig. 1c**). IF staining demonstrates precise spatial overlap of hCD59 and SYNPO within the glomerulus, further validating Pod-specific hCD59 expression (**SFig. 1c**).

To determine the dose regimen of ILY for ablating Pods, we conducted the terminal study with serial doses of ILY (200, 150, 75 and 37.5 ng/g body weight (BW), iv) to *ihCD59^+/-^/Nphs2Cre^+/-^* (experimental) and *ihCD59^+/-^/Nphs2Cre^-/-^* (control) mice. 75ng/g BW is the minimal dose of ILY to induce 100% of the mice death in *ihCD59^+/-^/Nphs2Cre^+/-^* (LD100). To ensure complete, specific, and acute ablation of Pods, we used 2.5X LD100 to characterize the consequences of Pod ablation in mice. IF staining of SYNPO (Pod-specific marker) and hCD59 shows no SYNPO or hCD59 signal in the glomeruli of experimental mice (**Fig. 1a**). In contrast, SYNPO signal is present in the glomeruli of ILY-injected control (**Fig. 1a**). Accordingly, 3hours after ILY injection (2.5X LD100), experimental but not control mice show reduced activity and signs of discomfort (**Supplemental Video 1**). After 6 hours, control mice still exhibit normal physical activity, whereas experimental mice show more pronounced reductions in activity and hunching (**Supplemental Video 2**). These results demonstrate that specific and complete Pod ablation only occurred in ILY-injected experimental mice. ILY injection caused hematuria (**Fig. 1b**) and albuminuria (**Fig. 1c, d, e, and SFig. 2**) within 6 hours, anuria within 6-24 hours, and death within 12-24 hours, along with elevated potassium, BUN, and creatine levels only in experimental mice (**Fig. 1f**). Histological analysis demonstrated that ILY injection at 2.5X LD100 mediated massive, global Pod damage and destruction of the glomerular tuft with widespread tubular proteinuria, degeneration, and necrosis only in experimental mice (**Fig. 1g**). Collectively, those results demonstrated that the specific ablation of Pods causes hematuria, albuminuria, progressive anuria, renal failure, and death.

**Figure. 1.**
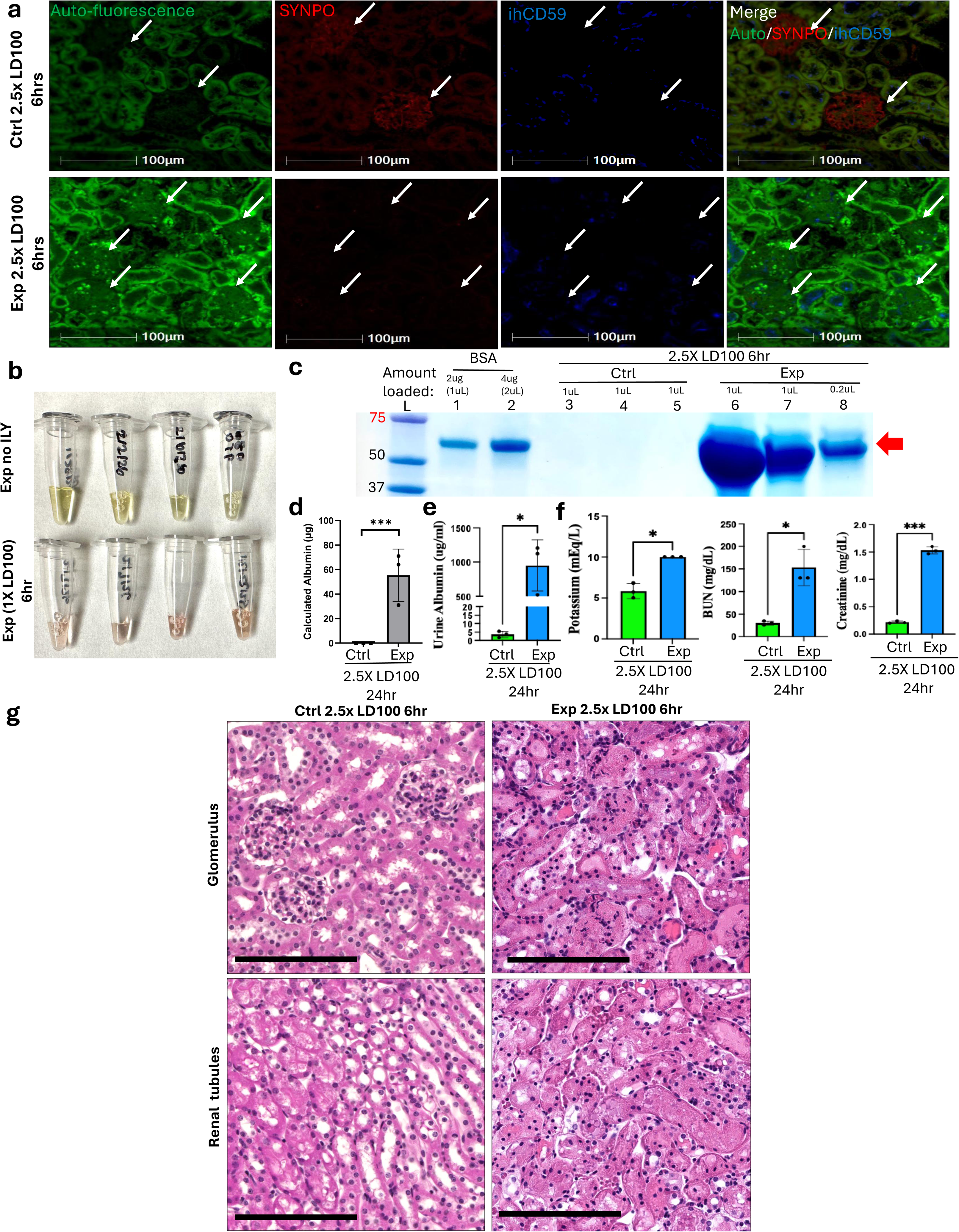
ILY-mediated Pod ablation induces glomerular disintegration and renal tubule necrosis (RTC) in *ihCD59⁺/⁻/Nphs2Cre⁺/⁻* mice: (a) Loss of podocyte-specific markers following ILY exposure. Immunofluorescence (IF) analysis confirmed ablation of hCD59⁺ podocytes in experimental (Exp, or *ihCD59^+/-^/Nphs2Cre^+/-^*) mice following ILY treatment (2.5 X LD100 at 6 hours (hrs), n=3). Complete loss of hCD59 (blue) and synaptopodin (red) signals within glomeruli (arrows) corresponded to destruction of the glomerular tuft. In contrast, littermate control (Ctrl, or *ihCD59^+/-^/Nphs2Cre^-/-^,* n=3) mice lacked hCD59 expression and retained intact podocyte structure and glomerular morphology (arrows). (b) Hematuria was present within 6 hrs post-ILY injection (2.5 X LD100) in Exp mice. (c) Visualization of albuminuria at 6 hrs post-ILY injection by SDS-PAGE, BSA: Bovine serum albumin.(d) Quantification of albumin level in c. (e) Quantification of urine albumin level in Ctrl and Exp mice(n=3) at 24 hrs post-ILY (2.5 X LD100). (f) Serum chemical analysis of BUN, potassium, and creatine levels in Ctrl and Exp mice at 24 hrs post-ILY injection. (g) Compared to controls (n=3), a 2.5X LD100 dose of ILY resulted in diffuse, global necrosis of glomeruli at 6 hrs post administration in the Exp mice (top right, arrows, n=3). Within the medulla, distal convoluted tubules and collecting ducts are diffusely filled with proteinaceous fluid and lined by RTCs in varying stages of degeneration and necrosis (bottom right). Bar=100um.

Ultrastructurally, transmission (TEM) and scanning electronic microscopic (SEM) analyses demonstrated the rapid loss of Pods in experimental mice resulted in expansion of glomerular fenestrations and filtration slits, loss of foot process, and partial denudation of the glomerular basement membrane (GBM) **(Fig. 2a, b, lower panel)** but not control mice **(Fig. 2a, b, upper panel).** Additionally, Pod-specific ablation resulted in loss of RTC brush border membranes (**Fig. 2c, lower middle panel**), mitochondria swelling (**Fig. 2c, lower right panel**), and enlarged RTC lumen (**Fig. 2d, lower panel**) in ILY-injected experimental but not control mice (**Fig. 2c, d, upper panel**). Taking together, these results demonstrated that specific and acute Pod destruction resulted in massive glomerular and tubular cell damage, leading to acute kidney injury.

**Figure 2:**
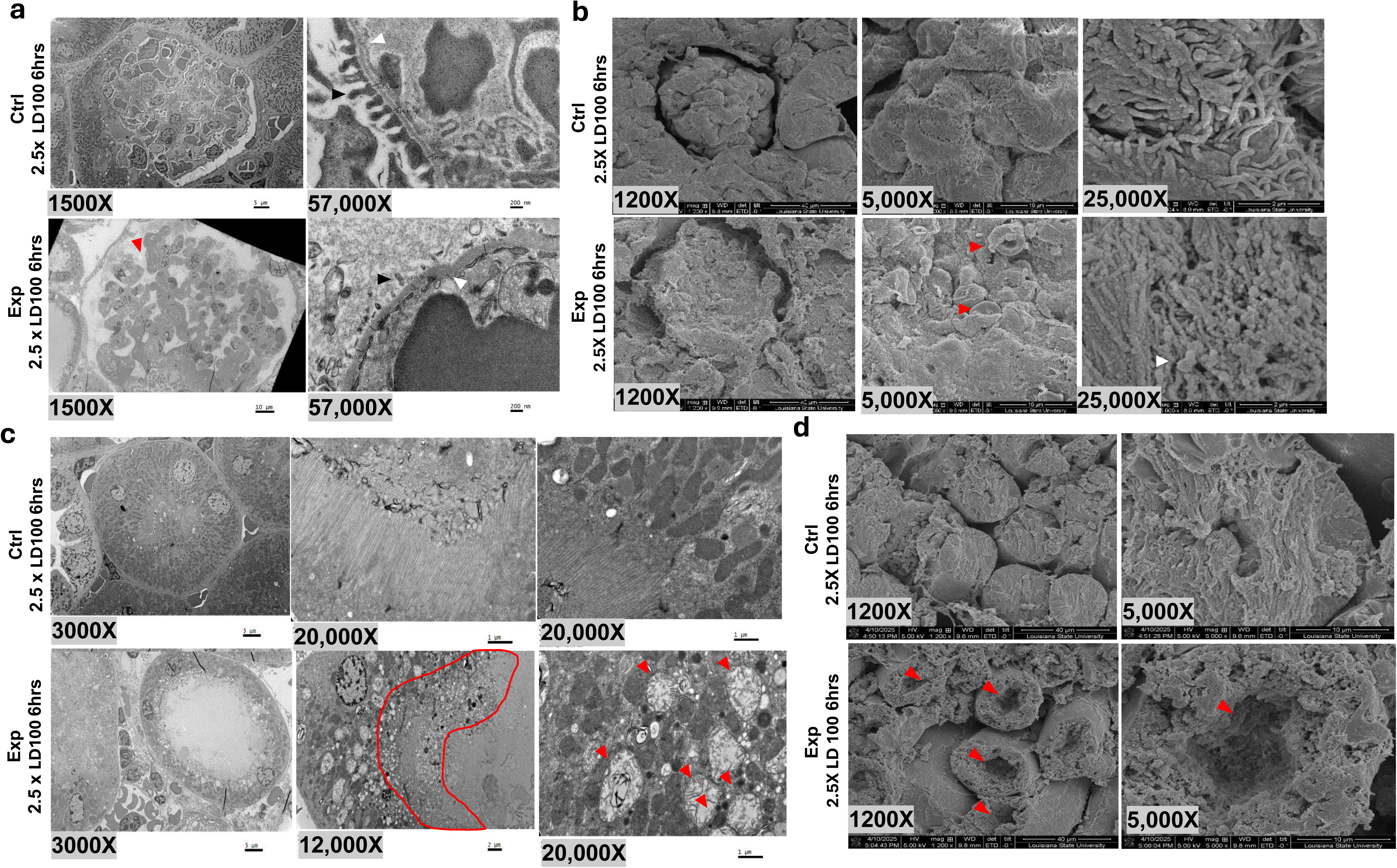
Ultrastructural studies show foot process damage in the glomerulus and mitochondria dysfunction in tubules 6 hr post-ILY injection in the mice: Transmission electron microscopy (TEM) of kidney tissue from experimental mice (*Exp, or ihCD59^+/-^ /Nphs2Cre^+/-^*,n=2) and control (Ctrl, or *ihCD59^+/-^/Nphs2Cre^-/-^,* n=2) mice 6 hours (hrs) after intraperitoneal ILY injection reveals extensive ultrastructural injury. (a) Low- (×1,500) and high-magnification (×57,000) TEM images demonstrate near-complete loss of podocytes (red arrow), with widespread effacement of foot processes (black arrow) and marked expansion of glomerular fenestrations and filtration slits (white arrows). (b) Scanning electron microscopy (SEM) of kidney tissue from experimental mice at 6 hrs post–ILY injection demonstrates extensive glomerular injury, including deformation and collapse of the renal corpuscle, effacement and malformation of podocyte foot processes (white arrow), and the aberrant presence of erythrocytes within Bowman’s space (red arrows) (c) TEM shows podocyte-specific ablation results in secondary proximal tubular injury, characterized by brush border loss (delineated by red line), cytoplasmic vacuolation, and intracellular lipid droplet accumulation. Mitochondria in tubular epithelial cells display pronounced swelling, indicative of early bioenergetic dysfunction. (d) Podocyte-specific ablation is accompanied by prominent cytoplasmic vacuolization and swelling in proximal tubular epithelial cells, with marked dilation of the tubular lumen (pointed by red arrows), consistent with early tubular injury and functional impairment.

### Dose-dependent effect on Pod ablation-mediated RTC necrosis and renal failure

To investigate whether the degree of Pod ablation correlates with the severity of renal failure associated with massive RTC damage, we conducted dose-dependent studies by injecting different doses of ILY (0.5X LD100, 1XLD100, and 2.5X LD100) in experimental animals. At 6 hours, doses higher than (2.5X and 1X) and less than LD100 (0.5X-0.75X) had 100% and 70% of Pod loss, respectively (**Fig. 3a, b**). ILY injection at doses higher than LD100 in experimental mice leads to loss of physical activity (**Fig. 3c**, **Supplemental Table 1**, and **Supplementary Video 3**) and shortening survival time post-injection, which correlated positively with ILY dose (**Fig. 3d**). Increased albuminuria level in mice is also positively correlated with the injected ILY doses (**Fig. 3e, f**). Consistently, histological analysis shows that nonlethal doses (0.5XLD100) resulted in minimal glomerular and tubular changes after 24h post-ILY administration, whereas lethal doses (LD100) of ILY resulted in acute, segmental to global, glomerular necrosis (arrow) and multifocal tubular necrosis by 6 hours post-administration (**Fig. 3g**). Supralethal dose (2.5X LD100) of ILY leads to diffuse, global glomerular necrosis (arrows) and subtotal tubular necrosis by 6 hours post administration (**Fig. 3g**). Increased levels of serum BUN and potassium (K) are also dependent on the injected ILY dose in experimental mice as compared to control mice (**Fig. 3h**). Of note, creatinine significantly rises at 1 X LD100, while high but not reaches significantly at 2.5 X LD100 due to the widespread data (**Fig. 3h**). Altogether, these results demonstrate that there is dose-dependent effect on Pod ablation-mediated tubular cell necrosis and renal failure.

**Figure 3.**
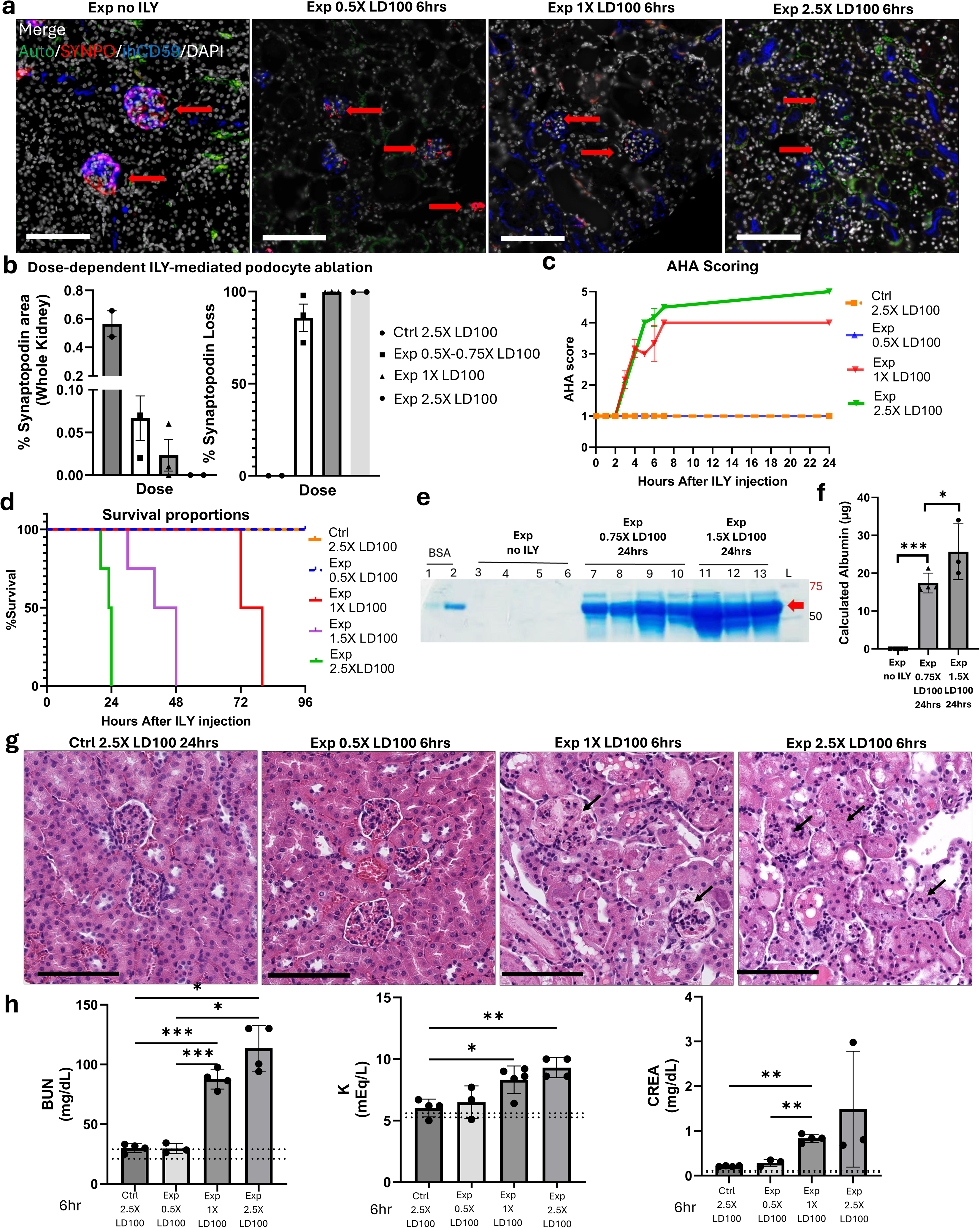
ILY mediated Podocyte depletion causes dose-dependent effect on survival rate, AHA behavior change, albuminuria, tubular cell damage, and renal failure in experimental mice: (a) Dose-dependent podocyte ablation in response to ILY exposure. Experimental mouse (*Exp, or ihCD59^+/-^/Nphs2Cre^+/-^*) glomeruli (red arrow) exhibit global expression of synaptopodin (red, SYNPO) and CD59 (blue). 0.5X LD100 6 hrs post-ILY (n=2) has segmental to global loss of SYNPO and CD59 in glomeruli (red arrows) of experimental mice. 1 X LD100 at 6 hrs-post injection in experimental mice (n=3) has a subtotal loss of SYNPO staining with glomerular tufts (red arrows). 2.5X LD100 at 6 hrs post injection experimental mice (n=3) glomeruli exhibit diffuse and global loss of SYNPO and CD59 (red arrow). White=DAPI, Green = Autofluorescence, Red = SYNPO, Blue = CD59. Bar = 100um. (b) % SYNPO+ area of whole kidney section (left panel) and % of podocyte loss (1-%glomerular SYNPO area in experimental mice divided by %glomerular SYNPO area in control mice). (c-d) ILY dose-dependent effect on animal health assessment (AHA) scoring (c), and survival rate (d). (e) Visualization of dose-dependent effect on albuminuria by SDS-PAGE. (f) Quantification of albumin level in e. (g) Dose-dependent ILY-mediated glomerular damage and tubular cell necrosis. Control mice have normal glomerular and tubular histomorphology. Experimental mice that received 0.5 X LD100 have minimal glomerular and tubular changes after 6 hrs post-ILY administration (n=2). 1 X LD100 dose of ILY results in diffuse, segmental to global glomerular necrosis (arrow) and multifocal tubular necrosis by 6 hrs post administration (n=3). 2.5XLD100 dose of ILY results in diffuse, global glomerular necrosis (arrows) and subtotal tubular necrosis by 6 hrs post administration (n=3). HE. Bar = 100 um (h) Measurement of BUN, K+, and Creatinine levels in serum of control and experimental mice at 6 hrs post-ILY injection (0.5XLD100, 1XLD100, and 2.5XLD100).

### Progressive development of renal failure and tubular damage in response to the initial Pod ablation

To explore whether the initial acute Pod damage directly and sequentially causes the rapid tubular cell necrosis, leading to renal failure and death, we measured serum chemicals for renal function, histologically analyzed the kidney at 3, 6 and 24-48 hours after ILY injection (1-1.5 X LD100) in the experimental mice. Serum BUN, potassium, and creatine levels were significantly higher at 6 or/and 48 hours than 3 hours post ILY injection (**Fig. 4a**). Histologically, the time-series studied revealed that dramatic protein deposition in tubule cell lumens (albumin casts) and Bowman’s space (at 3 hours) preceded proximal tubular necrosis which often manifested, even at lower ILY doses by 6 hours post ILY injection (**Fig. 4b**). After 24 hours post ILY injection, there is diffuse, global glomerular necrosis and subtotal tubular necrosis of both the proximal and distal tubules (**Fig. 4b**). These results clearly indicate that the initial acute Pod ablation caused protein accumulation in tubules first, and progressed over time to tubule cell necrosis, leading to renal failure and even death.

**Figure 4.**
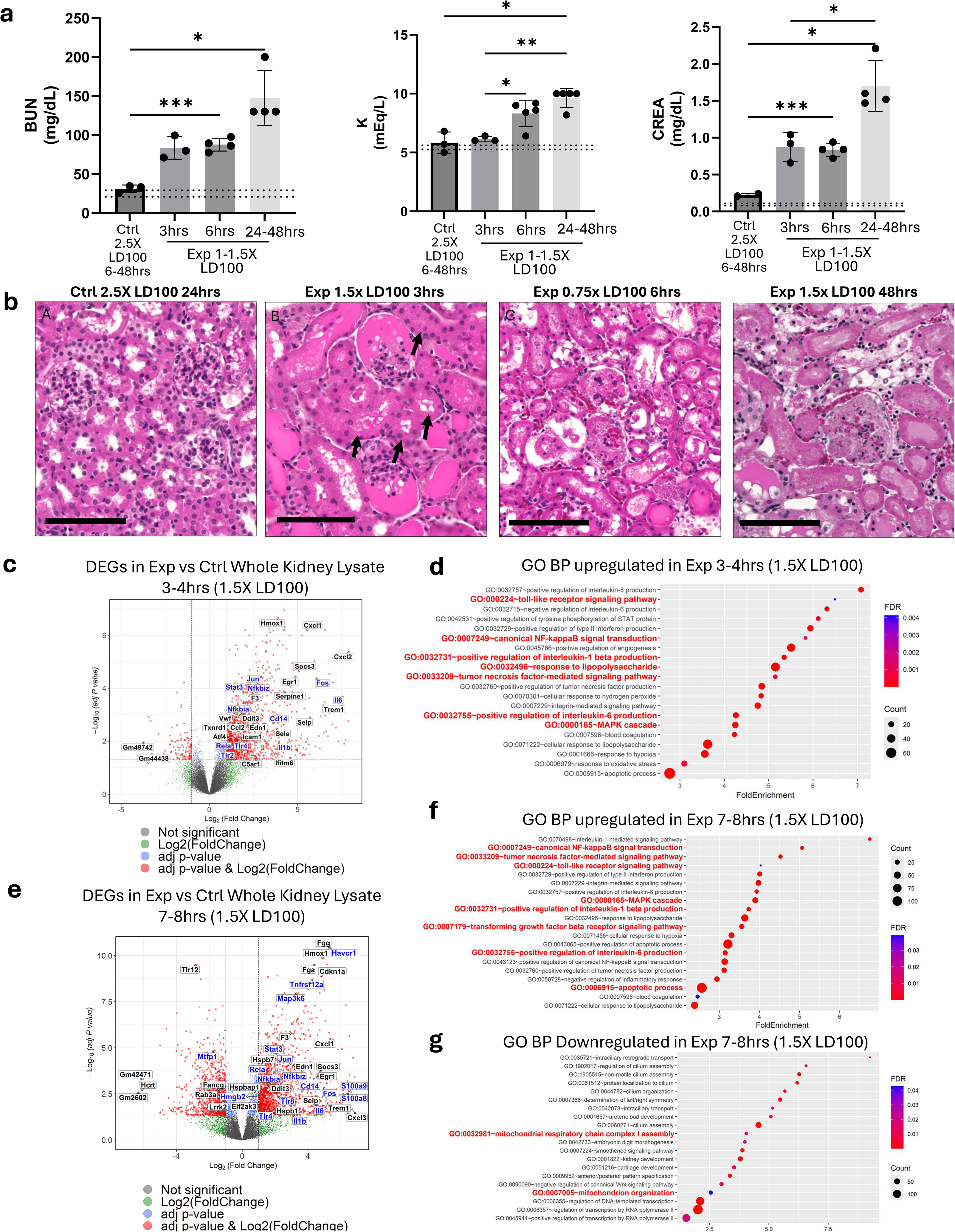
Time-dependent renal failure, tubular cell damage, and global transcriptomic changes in experimental mice: (a) Measurement of BUN, K+, and Creatinine levels in serum of control (Ctrl, or *ihCD59^+/-^/Nphs2Cre^-/-^)* and experimental (Exp, or ihCD59^+/-^ /Nphs2Cre^+/-^) mice at 3 hrs, 6 hrs, and 24-48 hrs post-ILY injection. (dashed lines = reference range) (b) In control mice absence of hCD59 spares kidneys from ILY-mediated podocyte ablation preserving normal renal histomorphology. At 3 hrs post-ILY injection, supralethal doses (1.5X LD100) result in severe proteinuria in Bowman’s spaces and renal tubules before tubular (arrows, viable proximal tubules) and glomerular necrosis. In contrast, by 6h post-ILY injection even sublethal doses (0.75X LD100) result in extensive necrosis of glomerular tufts and tubules. Animals that survive lethal doses for 48 h post-ILY (n=4) exhibit subtotal, global necrosis of glomeruli and renal tubules. (c) Volcano plot showing differentially expressed genes (DEGs) in the kidneys harvested 3-4 hr post-ILY treatment in experimental (n=4) versus controls (n=3). Y-axis shows −log10 (adjusted P value) (Benjamini–Hochberg). DEGs passing both cutoffs (padj <0.05 and |log2FC| >10)are displayed as significant (red). Tlr4 signaling associated genes in the significant DEGs are highlighted in blue. (d) Gene Ontology Biological Process over-representation analysis of upregulated DEGs in kidney lysate. (e) Volcano plot showing significant DEGs in whole kidney lysate *of ihCD59^+/-^ /Nphs2Cre^+/-^* mice 7–8 h post-ILY(n=3) with matched controls (n=3). Significant DEGs (padj<0.05 and |log2FC| >10) are associated with TLR–NF-κB/MAPK axis, inflammatory mediators, Kidney injury markers (Havcr1), and mitochondrial fusion protein (Mtfp1) (blue).(f-g) GOBP analysis of significantly up-regulated(f), and downregulated (g) DEGs in kidney lysate

To define early molecular pathways driving acute renal failure downstream of pod ablation, we performed bulk RNA-seq on whole-kidney lysates from control and experimental mice injected with ILY (1.5 LD100) and harvested at 3–4 h and 7–8 h post-injection (**Supplemental table 2**). Upregulated genes in 3-4h post-ILY experimental mouse kidney included genes related to TLRs (Tlr2, Tlr4, and Cd14), NF-κB signaling (Nfkbia, Nfkbiz, and Rela), JAK/STAT signaling (Stat3), MAPK signaling genes (Jun, Fos), and downstream proinflammatory cytokines (Il1b, Il6) (**Fig. 4c**). Significantly upregulated gene ontology biological process (GOBP) pathways in 3-4hr post-ILY ablated kidney lysate are related to responses to lipopolysaccharide (LPS), toll-like receptor signaling pathway, canonical NF-κB signal transduction, tumor necrosis factor (TNF)-mediates signaling pathway, MAPK cascade, positive regulation of interleukin-1 beta production, and positive regulation of interleukin-6 production (**Fig. 4d**). Due to few significantly downregulated DEGs, the only significantly downregulated pathway in 3-4hr post-ILY injected mice was related to regulation of DNA-templated transcription (**Supplementary table 3**). At 6-7hr post-ILY injection, TLR4-NFκB-MAPK signaling axis related genes and pathways continued to be significantly upregulated in experimental mice compared to WT mice (**Fig. 4e,f, SFig. 4a)**. Kidney injury markers Kim-1 (Havcr1), is significantly upregulated only at 6-7 hours post ILY injection, indicating initiation of acute kidney failure (**Fig. 4e**). Moreover, mitochondrial fusion related genes (Mtfp1), and mitochondrial related pathways are downregulated in experimental mice, implying initiation of mitochondrial dysfunction at 6-7hr post-ILY injection (**Fig. 4e, g, Sfig. 4b**), in-line with previous structural and ultrastructural changes in tubule cells as described above. Together, these results indicate that pod ablation directly contributes to progressive development of renal failure and tubular damage.

To further validate the changes of TLRs, mitochondrion genes and corresponding pathways as well as apoptotic pathways, we mined previously published transcriptomic datasets obtained from 3 different human glomerular dysfunction-related kidney disease cohorts, consisting of: 1) biopsied kidneys from advanced diabetes nephropathy patients, 2) glomerulus micro dissected from biopsied IgAN patient kidney, and 3) tubulointerstitium from biopsied IgAN, FSGS, LN, MN, and MCD patient kidneys (**Supplemental table 4**) (22–24). We plotted the log fold changes of the genes (**Fig. 5a-b**) and enrichment scores of the pathways (**Fig. 5c-h**) to directly compare their changes observed in our podocyte ablated mouse kidneys vs in the biopsied kidneys of the human cohorts (**Fig. 5**). To comprehensively reveal the dynamics of the pathway changes observed in our mouse study, we also included the enrichment scores of TLRs, mitochondrion function-related pathways and apoptotic pathways analyzed by Gene ontology (GO) and Gene Set Enrichment Analyses (GSEA) (**Fig. 5c-5h**). Consistently, we found significantly upregulated TLR2, TLR4, and/or TLR8 genes as well as TLRs and apoptotic pathways and downregulated mitochondrion function-related genes and pathways in glomerulus and/or tubulointerstitial tissues obtained from those 6 disease patients as compared with the respective controls (**Fig. 5a-g, Supplemental table 5, 6, 7, 8, 9**). Interestingly, before IgAN patients develop the profound kidney dysfunction, the tubulointerstitial tissues micro-dissected from the biopsied IgAN patient kidneys have significant up-regulation of TLRs gene and TLRs signaling pathways and down-regulation of mitochondrial signaling-related genes and pathways (24) than the micro-dissected IgAN glomerulus (23) (**Fig 5f-5h**). These profound changes were also observed in the tubuointerstitial tissues of FSGS, LN, MH and MCD (**Fig. 5f-5h**). These results indicate that glomerular dysfunction may trigger a more profound effect on those pathways in tubular cells as compared to that in glomerulus. The clinical data further demonstrated that glomerular dysfunction could increase TLR signaling activation and mitochondrial dysfunction, contributing to tubular cell damage.

**Figure 5.**
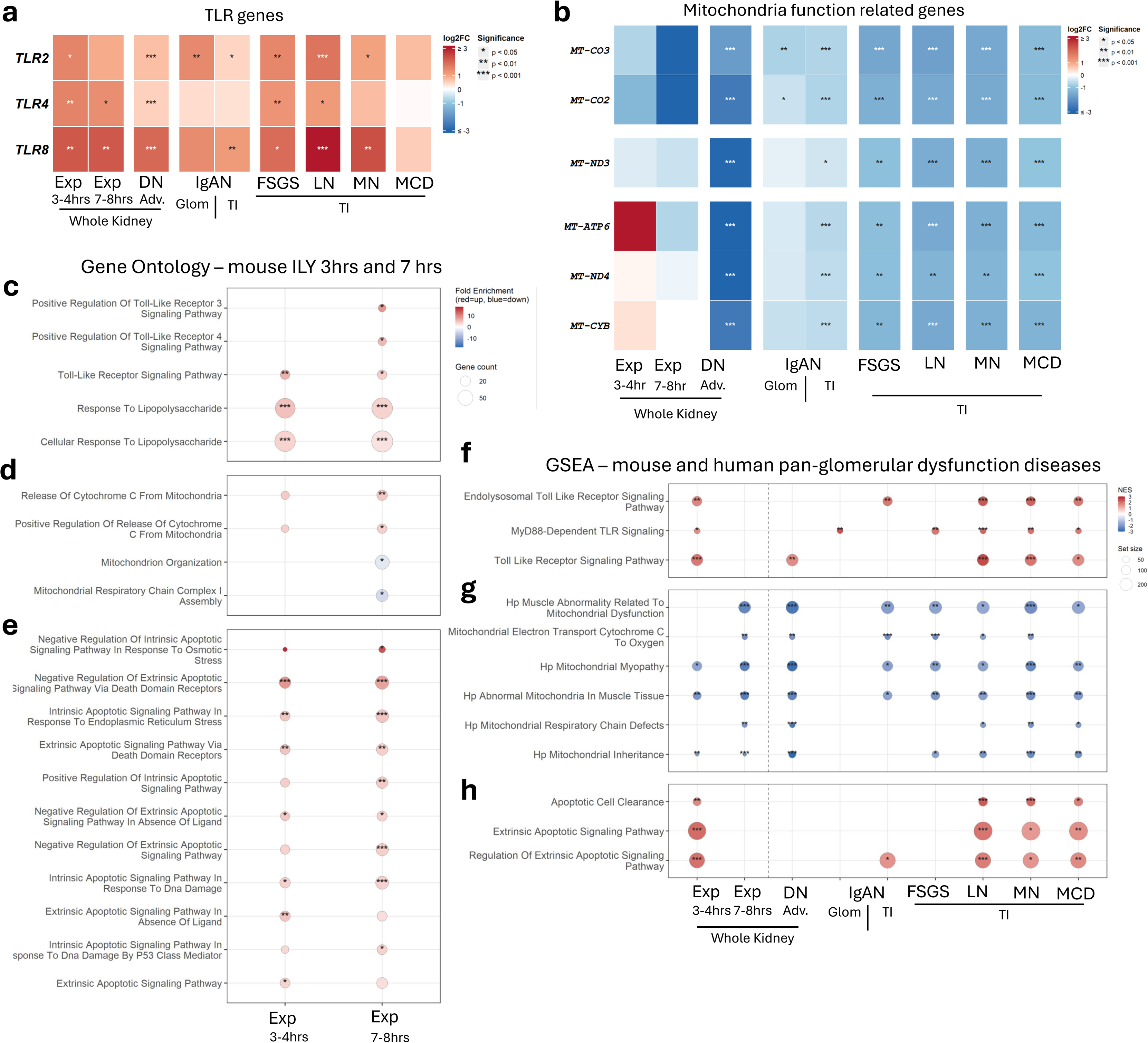
Conserved transcriptional changes of the TLR, mitochondrial function-related, and apoptotic signaling pathways in ablated-podocyte mouse and biopsied-glomerulonephritis patient kidney samples. (a-b) Heatmap showing log2 fold change of (a) TLR pathway genes (*TLR2*, *TLR4*, *TLR8*) and (b) mitochondria-related genes (*MT-CO3*, *MT-CO2*, *MT-ND3*, *MT-ATP6*, *MT-ND4*, *MT-CYB*) across mouse podocyte ablation (Exp 3-4hr and 7-8hr, whole kidney) and human glomerulonephritis datasets.(c–e) Bubble plots showing DAVID Gene Ontology Biological Process (GO BP) enrichment analysis in mouse podocyte ablation at 3-4hr and 7-8hr. Bubble color indicates fold enrichment (red = upregulated, blue = downregulated); bubble size represents gene count; asterisks indicate FDR significance. Pathways are grouped by category: (c) TLR/innate immune signaling, (d) mitochondrial function, and (e) apoptotic signaling pathways (intrinsic and extrinsic). (f–h) Bubble plots showing Gene Set Enrichment Analysis (GSEA) across murine podocyte ablation and human glomerular disease datasets. Bubble color indicates normalized enrichment score (NES; red = positive, blue = negative); bubble size represents gene set size. Pathways are grouped into the following categories: (f) TLR/innate immune signaling, (g) mitochondrial dysfunction, and (h) apoptotic signaling pathways. Asterisks indicate statistical significance (* p < 0.05, ** p < 0.01, *** p < 0.001). Abbreviations: DN: Diabetes Nephropathy; IgAN: IgA Nephropathy; FSGS: Focal Segmental Glomerulosclerosis; LN: Lupus Nephritis; MN: Membrane Nephropathy; MCD: Minimal Changes Disease. Glom: glomerulus; TI: Tubulointerstitial tissue.

### Acute specific Pod ablation resulted in severe glomerular sclerosis, ongoing tubular cell degeneration, and interstitial fibrosis

To explore the role of acute Pod damage on chronic kidney disease, we used lethal and non-lethal doses of ILY to inject into the experimental mice. After lethal dose injection, we also conducted peritoneal dialysis (PD) in the experimental mice treated with ILY as described(20) to extend the survival of the mice beyond 6 six weeks after ILY injection. PD was started at 6 hours post-ILY injection (1.5 LD100), at which time control mice (*ihCD59^+/-^/Nphs2Cre^-/-^*) showed a normal level of activity, while experimental mice showed no physical response to stimulus (**Supplemental Video 4**). After 2 h with PD treatment (8 hours post-ILY injection) the experimental mice partially restored normal activity with increased appetite (**Supplemental Video 5**). At 24 hours post-ILY injection, the experimental mice restored physical response to stimulus under the PD treatment. The mice recovered thereafter with normal physical activity with PD (**Supplemental Video 6**). Of the two PD mice, one of the animals was harvested at 2 weeks, and the other mouse was harvested at 6 weeks. We analyzed the urine albumin levels weekly and found increased albuminuria after ILY injection, which increased from week 1 to week 2, dramatically reduced from week 2 to week 3, and stabilized levels in week 4-6 as compared with the control (**Fig. 6a-e, SFig. 5a-b**). We also found that immunoglobulin (IgG) levels are highest at week 1, second highest in week 2, and gradually disappear from week 3 to 6 (**Fig. 6f-g, SFig. 5c**). Consistently, body weight was reduced by 13% at week 1 and evidently regained after week 2 post injection (**SFig. 6**). These results indicate that 2 weeks after Pod ablation induced AKI, there is partial, spontaneous amelioration of glomerular and tubule injury. Despite partial recovery, the PD mice developed multifocal glomerulosclerosis (**Fig. 6e, SFig. 7a-g**), ongoing tubular degeneration (arrows in **SFig. 7d-f**). PAMS staining shows that in 6 weeks post-PD mice, there is increased thickness of tubular basement membranes in regions with tubular loss and interstitial fibrosis (**SFig. 7h-i**).

**Figure 6.**
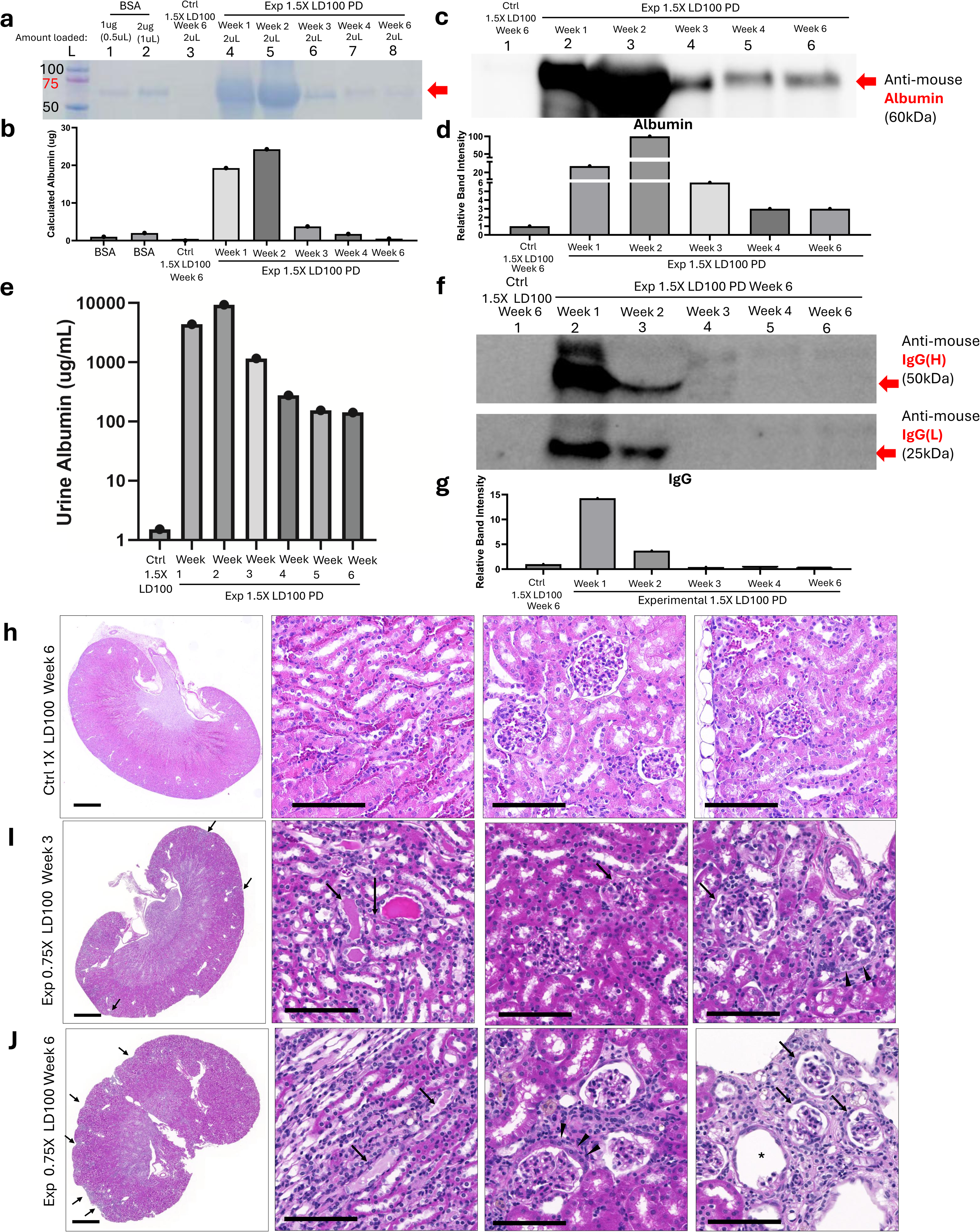
Specific podocyte ablation results in chronic progressive renal glomerular sclerosis and interstitial fibrosis despite partial restoration of function. (a) Visualization of proteinuria by SDS-PAGE from week 1 to 6 in the experimental mouse (Exp, or *ihCD59^+/-^ /Nphs2Cre^+/-^*) that received fluid exchanges via PD catheter for 4 times/hours for 4 hours daily for two days after 6 hours (hrs) post-ILY injection (1.5 XLD100). BSA: Bovine serum albumin. (b) Quantification of calculated total protein level shown in Fig. 6a. (c) Western blot showing albumin level in urine from week 1 to 6 in that the PD-treated experimental mice. (d) Quantification of the relative albumin band intensity in the western blot in c. (e) Quantification of urine albumin level in control and in week 1-6 of the PD treated experimental mouse. (f) Decreasing infiltrated immunoglobulin (IgG) levels visualized by western blot in week 1-6 of the PD treatment mouse. (g) Quantification of the relative immunoglobulin band in western blot in Fig. 6f. (h) Control mouse showing a regular contour to the capsular surface of the kidney (Left), proximal and distal tubules (Middle right), and glomeruli (Middle right & Right) with normal histomorphology. (i) Sublethal ILY dose (0.75X LD100) 3 weeks post administration (n=3). (Left) The kidney’s contour is irregular due to multifocal regions of fibrosis and stromal collapse (arrows). Bar = 1mm. (Middle left) There is persistent proteinuria and tubular degeneration of renal tubules (arrows). (Middle right) Glomeruli are undergoing sclerosis (arrow). (Right) In regions of stromal collapse, glomeruli form synechiae (arrow) and Bowman’s capsule is lined by hypertrophic PECs (arrowhead). Bar = 100um. (j) Sublethal ILY dose (0.75X LD100) 6 weeks post administration (n=2). (Left) The contour of the kidney is irregular due to multifocal regions of fibrosis and stromal collapse (arrows). Bar = 1 mm. (Middle left) There is ongoing proteinuria and tubular degeneration within distal segments of the nephron (arrows). (Middle right) Bowman’s capsules surrounding remaining glomerular tufts are often lined by hypertrophic PECs (arrowheads). (Right) Glomeruli within regions of fibrosis are often sclerotic (arrows) or absent (asterisks). Bar = 100 um.

Similarly, we also used a non-lethal dose of ILY (0.75X LD100) to inject experimental mice and harvested the mice at 3 weeks and 6 weeks. At both 3 and 6 weeks post ILY injection, experimental mice have multifocal regions of fibrosis and stromal collapse, persistent proteinuria, tubular degeneration, and ongoing glomerulosclerosis (**Fig. 6h-j**). In regions of stromal collapse, glomeruli form synechiae/crescents and Bowman’s capsule is often lined by hypertrophic parietal epithelial cells (PECs) (**Fig. 6h-j**). Consistently, the albumin and immunoglobulin levels in the urine collected from one ILY non-lethal dose injected-experimental mouse was dramatically increased at week 1 then dramatically decreased at week 2 post injection (**SFig. 8**), a similar changed patten as seen in PD mice received with 1.5 LD100 dose of ILY. Together, those results demonstrate that despite partial recovery of glomerular and renal function, acute Pod ablation leads to progressive, irreversible glomerular sclerosis and interstitial fibrosis with ongoing tubular cell degeneration.

## Discussion

Here, we report on the establishment and characterization of acute, specific and conditional Pod ablation mouse model via ILY-hCD59 cell ablation tool and document that the Pod ablation causes hematuria, albuminuria, progressive anuria, renal failure and even death in mice in a dose dependent manner. These results are comparable with the previously published findings obtained by widely used, nonspecific, and slower acting cell ablation models (LPS, chemicals, murine knockout models or diphtheria toxin [DT]/DT receptor [DTR])(4, 17), and other toxin-induced models of Pod damage(1, 2, 5–13)). Compared with these tools, the ILY/ihCD59 cell-ablation method has several attractive features, including high specificity without any detectable off target effects, a wide pharmacological window, and a rapid ablation mechanism independent of DNA replication or protein synthesis(14, 16). DT-mediated Pod damage causes renal failure in mice within 5-21 days (3) and in rats within 10-20 days after DT injection (1), which is due to DT-mediated slow killing effect on Pods. Here, we report that ILY injection with lethal doses results in the complete ablation of hCD59 expressing Pods specifically in the experimental mice leading to hematuria, albuminuria, progressive anuria, and renal failure in the mice in 6 hours or less, with progression to death within 1-3 days. The rapid development of the renal failure and death was also observed in a mouse model for congenital nephrotic syndrome by targeted deletion of the nephrin gene (*Nphs1*) in mice (25). Homozygous *Nphs1* knockout mice *(Nphs1^-/-^*) born at an expected frequency of 25% had a normal activity at birth but immediately developed massive proteinuria and edema and died within 24 h(25). The kidneys of *Nphs1^-/-^* mice exhibited enlarged Bowman’s spaces, dilated tubules, effacement of Pod foot processes and absence of the slit diaphragm(25). Similarly, human NPHS1 patients develop proteinuria in utero and usually develop nephrotic syndrome, with massive non-selective proteinuria and edema, and early death without renal transplantation following extensive medical care(25–27). The progressive development of massive proteinuria and structural changes in NPHS1 patients is similar to what we observed in our Pod cell ablation model. Taken together, the acute, specific and conditional Pod ablation mouse model reported here provides another new tool to investigate the pathophysiological role of Pod cells in the pathogenesis of Pod damage-related human diseases such as nephrotic syndrome (NS), autoimmune related nephropathy (ARN), and diabetic nephropathy (DN).

We utilized the ILY-mediated hCD59 expressing Pod ablation model in a time-series study to document that the rapid loss of Pods and foot processes results in massive protein accumulation in the lumen of RTCs first, prior to RTC damage and acute RTC necrosis, which progresses to renal failure and even death. Further, ILY-treated mice that survived beyond 6 weeks, either by treating mice given a lethal dose of ILY with peritoneal dialysis (PD) or by given mice a non-lethal dose of ILY injection, had persistent albuminuria, glomerulosclerosis, ongoing RTC degeneration, and extensive interstitial fibrosis. These results provide direct experimental evidence supporting the notion that Pod damage causes acute RTC necrosis and interstitial fibrosis. Furthermore, we conducted the bulk RNA analysis of the kidneys after ILY injection to mice. The analysis revealed upregulated Tlr genes (Tlr 2, 4, and 8), Tlr signaling pathways, and necrotic and apoptotic pathways, and downregulated mitochondrial function-related genes in pod ablation, which is further validated by the transcriptomic analyses of 3 different human glomerular dysfunction-related kidney cohorts(22–24). Consistently, the filtration of high molecular weight blood proteins, including albumin, immune globulins, complement proteins, etc., through a glomerular barrier compromised by Pod damage has long been implicated in causing proximal tubule injury and necrosis(28–32). Supportively, Dr. Batuman’s group has also found that aristolochic acid I and antibody light chains induce tubule injury through the activation of TLRs such as TLR4(31, 32). These findings are consistent with other findings showing that 1) TLR4 promotes interstitial fibrosis but attenuates tubular damage in unilateral ureteral obstruction mouse model(33), and 2) TLR4 signaling in renal tubular epithelial cells has also been implicated in the mediation of inflammation and tissue Injury in AKI(34–36) and a murine kidney transplant model(37). These results, together with our observation regarding TLRs-signaling activation induced by Pod ablation, support the role of TLRs in the pathogenesis of AKI, which warrants further investigation with our specific, acute, and conditional Pod ablation model.

Additionally, we observe a dramatic recovery of the broken glomerular infiltration barrier within 2 weeks in response to acute and specific Pod ablation. Proteinuria is a clinical hallmark of kidney disease(38). Selective albuminuria is a critical, early biomarker of glomerular injury, specifically associated with minimal change nephrotic syndrome (MCNS) and Pod dysfunction(38–40). Non-selective albuminuria involves leakage of both low and high-molecular-weight proteins (e.g., IgG, IgM) due to significant structural damage, often indicating severe conditions like focal segmental glomerulosclerosis or advanced nephropathy(38–40). Consistently, we observed that the mice had hematuria often occurring at first 24 hours, albuminuria that was extremely high in week 1, peaked at week 2, and dramatically decreased but remained high at weeks 3-6. Immunoglobinuria was similar but with more rapid kinetics, peaking at week 1, dramatically decreasing but remaining elevated at week 2, and disappearing between weeks 3-6 after the Pod ablation with either non-lethal or lethal dose of ILY followed by PD to extend the mice survival. The presence of red blood cells and increased level of immunoglobulins in urine, and albuminuria together with massive glomerular destruction in these mice at the acute phase (6 hours to 7 days) post Pod ablation clearly indicates that the specific Pod ablation leads to more advanced glomerular injury such as dysfunction of glomerular base membrane, endothelial cells, and even mesangial cells, which warrants further investigation. Furthermore, the change of non-selective albuminuria in the mice at week 1 to selective albuminuria at week 2 post Pod ablation suggests at least partial restoration of the broken glomerular filtration barrier. Further understanding of the detailed pathological changes, and molecular and cellular mechanisms underlying Pod ablation-mediated glomerular disruption and repair will advance our understanding of Pod biology and develop efficient targeted treatments for kidney diseases.

## Methods

### Animal model strains and their maintenance

*Nphs2Cre* hemizygous (*Nphs2Cre^+/-^*) mice (JAX 008205) were obtained from The Jackson Laboratory (Bar Harbor, ME) and housed at the Tulane University School of Medicine. The *ihCD59* homozygous *(ihCD59^+/+^*) mice (JAX 035504), previously generated and backcrossed onto a C57BL/6 genetic background for at least seven generations, were used for breeding. To generate experimental genotypes, *Nphs2Cre^+/-^* mice were crossed with *ihCD59^+/+^* mice to produce the following offspring: *ihCD59^+/−^/Nphs2Cre^+/-^*(experimental mice (Exp)) and *ihCD59^+/−^/Nphs2Cre^-/-^* (control mice (Ctrl)). All animal experiments were conducted in accordance with the ethical guidelines of the Institutional Animal Care and Use Committee (IACUC) at Tulane University (Protocol number 2335). Mice were housed in a specific pathogen-free (SPF) facility at the Tulane University School of Medicine, maintained under a 12-hour light/dark cycle with controlled environmental conditions.

### Intermedilysin purification

Recombinant His-tagged ILY was purified using the HisBind Purification Kit (EMD 70239) following the manufacturer’s instructions, as previously described (15, 16). Briefly, bacterial cultures expressing His-tagged ILY were lysed, and the protein was captured using a nickel-affinity chromatography column. After washing to remove non-specifically bound proteins, the His-tagged ILY was eluted under optimized conditions. The concentration and purity of the purified ILY were assessed by SDS-PAGE, ensuring adequate purity for downstream applications.

### Podocyte depletion

Age-matched *ihCD59^+/−^/Nphs2Cre^+/-^ (Exp)* and *ihCD59^+/−^/Nphs2Cre^-/-^ (Ctrl)* mice were injected with various doses (lethal and non-lethal doses) of ILY (37.5-400 ng/g body weight) via tail vein (i.v.). In total, the mice were euthanized at different time points to see the acute and chronic effects of ILY after podocytes ablation. Kidney was harvested after blood perfusion, blood was collected and spot urine was collected before and after ILY administration at different time intervals.

### Peritoneal Dialysis

Peritoneal Dialysis catheter insertion and peritoneal dialysis were performed as described(20). Briefly, at 8–10 weeks of age, mice underwent surgical placement of a customized peritoneal dialysis (PD) catheter into the peritoneal cavity as previously described(20). After 7 days of postoperative recovery, rapid ablation of hCD59-expressing podocytes was induced by intravenous administration of intermedilysin (ILY) at 1.5 × LD100. Beginning 6 h after ILY injection, mice (n=2) received PD fluid exchanges via the indwelling catheter with a low-calcium peritoneal dialysis solution (2.5 mEq/L Ca²⁺; Baxter, L5B4825). PD was performed at a rate of 4 exchanges per hour for 4 h daily for 2 consecutive days, with 1.5 mL of dialysate administered per exchange. Mice were then followed longitudinally and harvested in week 6 after ILY-induced injury.

### Urinary Albumin Quantification by ELISA

Albuminuria, a hallmark of glomerular barrier disruption, was quantified using a commercially available rat-specific Albumin ELISA Kit (e.g., Abcam, USA) following the manufacturer’s instructions. Briefly, urine samples were thawed on ice and centrifuged at 10,000 g for 5 min to remove cellular debris. Samples were diluted (typically 1:2000 to 1:256000) in assay buffer to ensure readings fell within the linear range of the standard curve. Absorbance was measured at 450 nm using a microplate reader.

### Urine Protein Fractionation by SDS-PAGE

To qualitatively assess the molecular weight distribution of excreted proteins and confirm albuminuria, sodium dodecyl sulfate-polyacrylamide gel electrophoresis (SDS-PAGE) was performed according to the method of Laemmli. Urine samples were mixed with 4× Laemmli sample buffer containing 10% beta-mercaptoethanol and denatured at 100°C for 5 minutes. Equal volumes of urine (1 μL) were loaded onto 4–20% gradient polyacrylamide gels (Bio-Red) to capture a wide range of molecular weights. Electrophoresis was performed at a constant 95V for approximately 90 minutes. Gels were stained with Coomassie Brilliant Blue R-250 (0.1% w/v in 10% acetic acid and 40% methanol) for 1 hour, fixed with (10% Acetic acis+40% Methanol), and de-stained until distinct bands were visible. The presence of a dominant ∼66 kDa band indicated albuminuria (glomerular injury). Band intensity was quantified using ImageJ. Protein concentration for each sample was determined by normalizing sample band intensity to the BSA control and multiplying by the known concentration of the BSA standard.

### Serum Chemistry

After euthanasia, serum was collected from mice, and clinical chemistry analyses were performed using a Beckman AU480 chemistry analyzer at the Tulane National Primate Research Center (TNPRC) Clinical Pathology Core (RRID: SCR_024609).

### Immunohistochemistry Method

4 um tissues sections are mounted on Superfrost Plus Microscope slides, baked for 3 hours at 60°C and passed through Xylene, graded ethanol, and double distilled water to remove paraffin and rehydrate tissue sections. A microwave is used for heat-induced epitope retrieval. Slides are boiled for 20 minutes in a Tris based solution, pH 9 (Vector Labs H-3301), containing 0.1% Tween20. Slides are briefly rinsed in hot, deionized water and transferred to a hot citrate-based solution, pH 6.0 (Vector Labs H-3300) where they were allowed to cool to room temperature. Slides are removed from the antigen retrieval solution, washed in phosphate-buffered saline, and counterstained with Dapi. The slides are then washed in phosphate-buffered saline, deionized water, and Roche reaction buffer before being loaded on the Ventana Discovery Ultra autostainer where they would undergo sequential rounds of blocking, primary antibody incubation, washing, secondary antibody incubation, washing, and color development (**Supplemental Table 10**). Roche protocols for denaturation and neutralization are performed between rounds of staining to remove the first antibody and quench any remaining horse radish peroxidase. A second counterstain with Dapi is done on the machine. Upon removal, slides undergo alternating manual washes of deionized water containing 0.1% Dawn dish soap and plain deionized water for a total of 5 cycles before being permanently mounted. After drying overnight, slides are digitally imaged at 20X with a Zeiss Axio Scan.Z1. Image analysis was performed by a board-certified, veterinary pathologist using HALO HighPlex FL v4.1.3 from Indica Labs.

### H&E method

Tissue samples were collected in Zinc formalin (Anatech) and fixed for a minimum of 24 hours before being washed and dehydrated using a Thermo Excelsior AS processor. Upon removal from the processor, tissues were transferred to a Thermo Shandon Histocentre 3 embedding station where they were submersed in warm paraffin and allowed to cool into blocks. From these blocks, 4um sections were cut and mounted on charged glass slides, baked overnight at 60oC and passed through Xylene, graded ethanol, and double distilled water to remove paraffin and rehydrate tissue sections. A Leica Autostainer XL was used to complete the deparaffinization, rehydration and routine hematoxylin and eosin stain. Slides were digitally imaged with a Zeiss Axio Scan.Z1 and subsequently examined by a board-certified, veterinary pathologist using HALO software (Indica Labs).

### Western blot analysis

Urea samples were diluted in PBS and mixed with 4× Laemmli SDS sample buffer (Bio-Rad). Denatured proteins were resolved on 4-20% Criterion™ TGX™ precast midi protein gel (Bio-Rad) and transferred onto PVDF membranes using the Bio-Rad Trans-Blot turbo transfer system. Membranes were blocked with EveryBlot blocking buffer (Bio-Rad) and incubated with Horse Anti-mouse IgG, HRP-linked antibody (#7076, Cell Signaling, 1:5000) for 1 h at room temperature to detect immunoglobulins. Membranes were washed four times with TBS containing 0.1% Tween-20 and IgG was visualized using SuperSignal™ West Atto ultimate sensitivity substrate (ThermoFisher Scientific). Chemiluminescent signals were captured with a ChemiDoc™ MP imaging systems (Bio-Rad). Following detection, membranes were then stripped using Thermo Scientific restore stripping buffer, reblocked and incubated with mouse albumin polyclonal antibody conjugated with HRP (A90-134P, ThermoFisher Scientific; 1:4000) for 1 h at room temperature. After washing with TBST, albumin was visualized using SuperSignal™ West Pico PLUS chemiluminescent substrate (Fisher Scientific).Band intensity was quantified using ImageJ. Relative band intensity was calculated by normalizing the sample band intensity to that of the control.

### Bulk RNA-seq sample harvesting and total RNA isolation

12-week-old control (n=3) and experimental mice (n=6) were injected with a lethal dose of ILY. Those mice were harvested at either 3-4 hours or 7-8 hours after ILY injection. Kidney tissues were collected in 1 mL Trizol reagent (15596026; Invitrogen) and extracted with RNeasy Mini Kit (Cat. No.74104; QIAGEN, Hilden, Germany) following the manufacturer’s protocol. The concentration of RNA was determined by NanoDrop 2000.

### Bulk RNA-sequencing (RNA-seq) and Data analysis

Library preparation, total RNA-sequencing were performed at Tulane Center for Translational Research in Infection & Inflammation NextGen Sequencing Core as described previously (21). Downstream analysis, including raw read QC, alignment, abundance estimation, and gene-level differential expression, was also performed as described previously (21). Briefly, Raw reads were quality-checked with FastQC, aligned to mm10 using STAR, and quantified with featureCounts. Differential gene expression analysis was performed with DESeq2. Genes with adjusted *p* < 0.05 and |log2 fold change| > 1 were considered significantly differentially expressed.

All data handling, filtering, and figure generation were performed in R, primarily using the packages openxlsx, dplyr, tidyr, stringr, ggplot2, ComplexHeatmap, and circlize. Volcano plots were generated using the Enhanced Volcano package (v1.16.0) to visualize gene-level changes. Gene Ontology Analysis was conducted using the DAVID Gene Ontology server. Gene set enrichment analysis (GSEA) was conducted using r package clusterProfiler (v4.14.6). Mouse-specific gene sets from the C2 (curated) and C5 (Gene Ontology) categories of the MSigDB database were retrieved using msigdbr (v7.5.1). Gene sets with adjusted p-value < 0.05 were considered significantly enriched.

### Analyzing publicly available transcriptomic datasets of 3 different human glomerular dysfunction-related kidney disease cohorts and comparing them with our mouse datasets

Public human bulk RNA seq original datasets were obtained from GEO, which consist of 1) GSE142025 from whole kidney biopsied from advanced diabetic nephropathy patients (DN_adv), 2) GSE141295 of micro dissected glomerular samples from biopsied IgA nephropathy kidneys (IgAN_Glom), and 3) GSE175759 of micro dissected tubulointerstitium from biopsied IgAN (IgAN_TI), focal segmental glomerulosclerosis (FSGS_TI), lupus nephritis (LN_TI)), membranous nephropathy (MN_TI), and minimal change disease (MCD_TI) kidneys (**Supplemental table 4**). We performed the DEG and GSEA analysis of each disease cohort by comparing it with the corresponding control group as described above. For the cross-dataset comparisons, representative TLR-related genes (TLR2, TLR4, and TLR8) and representative mitochondrial function-related genes (MT-CO3, MT-CO2, MT-ND3, MT-ATP6, MT-ND4, and MT-CYB) log fold changes were extracted from the DESeq2 results and visualized as heatmaps. Gene symbols between human and mouse across datasets are converted using the R package msigDB (v7.5.1).. To identify pathways that were directionally consistent across our murine podocyte-ablation model and human kidney disease cohorts, GSEA outputs were screened using keyword-based pathway classification. To reveal the significantly conserved-up (NES > 0) or down regulation (NES < 0) of the pathways, q < 0.05) across multiple groups in at least one murine dataset and at least three clinical datasets, we plotted TLR-related and apoptosis-related, and mitochondrial pathways respectively.

### Statistical Analysis

Data are presented as mean ± standard error of the mean (SEM). For comparisons between two groups, unpaired Student’s *t*-tests were used. For experiments involving multiple groups and time points, two-way ANOVA, and mixed models with Tukey’s multiple comparisons were used. A *P* value < 0.05 was considered statistically significant. All analyses were performed using GraphPad Prism version 10.1.

## AUTHOR CONTRIBUTIONS

Conceptualization: X.Q., V.B, R.B, RVB. Methodology: YC., M.I., X.D., J. E., S.L., C.M., R.S., JLZ, JKK, R.B., and X.Q. Investigation: YC., M.I., X.D., J. E., A.S., G.M.H., S.L., C.M., R.S., JLZ, JKK, R.B., and X.Q., Data analysis and result interpretation: Y.C., M.I., V.B, R.B, RVB, X.Q. Funding acquisition: X.Q. Project administration: X.Q. Y.C. M.I. Writing – review & editing: X.Q. Y.C., M.I. JKK, V.B.,R.B, RVB.

## Data and materials availability

All data associated with this study are present in the paper or the Supplementary Materials. The data that support the findings of this study are available from the corresponding author upon reasonable request. The datasets generated and analyzed during the current study are available from the corresponding author and first author upon reasonable request. Wild-type C57BL/6 and *Nphs2Cre^+/-^*are available in the Jackson Laboratory (Bar Harbor, ME). The *ihCD59^+/+^* mice, previously generated and backcrossed onto a C57BL/6 genetic background for at least seven generations is available in the Jackson Laboratory (Bar Harbor, ME). ILY are available upon request through the material transfer agreements (MTA). Bulk RNA sequencing data reported in this paper will be deposited to the GEO database after the acceptance and will be available publicly at the date of publication (accession number: pending).

## Ethical approval

Our research follows Tulane University’s Institutional Animal Care and Use Committee-approved protocols (2335) and complied with all relevant ethical regulations for animal treatment.

## Acknowledgements

We thank Tulane National Biomedical Research Center P51OD011104 RRID: SCR_008167 Anatomic Pathology Core; RRID: SCR_024606; and Confocal Microscopy and Molecular Pathology Core: RRID: SCR_024613; TNBRC Clinical Pathology Core Facility RRID: SCR_024609

