## Supplemental Figures and Tables for "Rapid Podocyte ablation Causes Acute Renal Tubule Cell Necrosis and Interstitial Fibrosis"

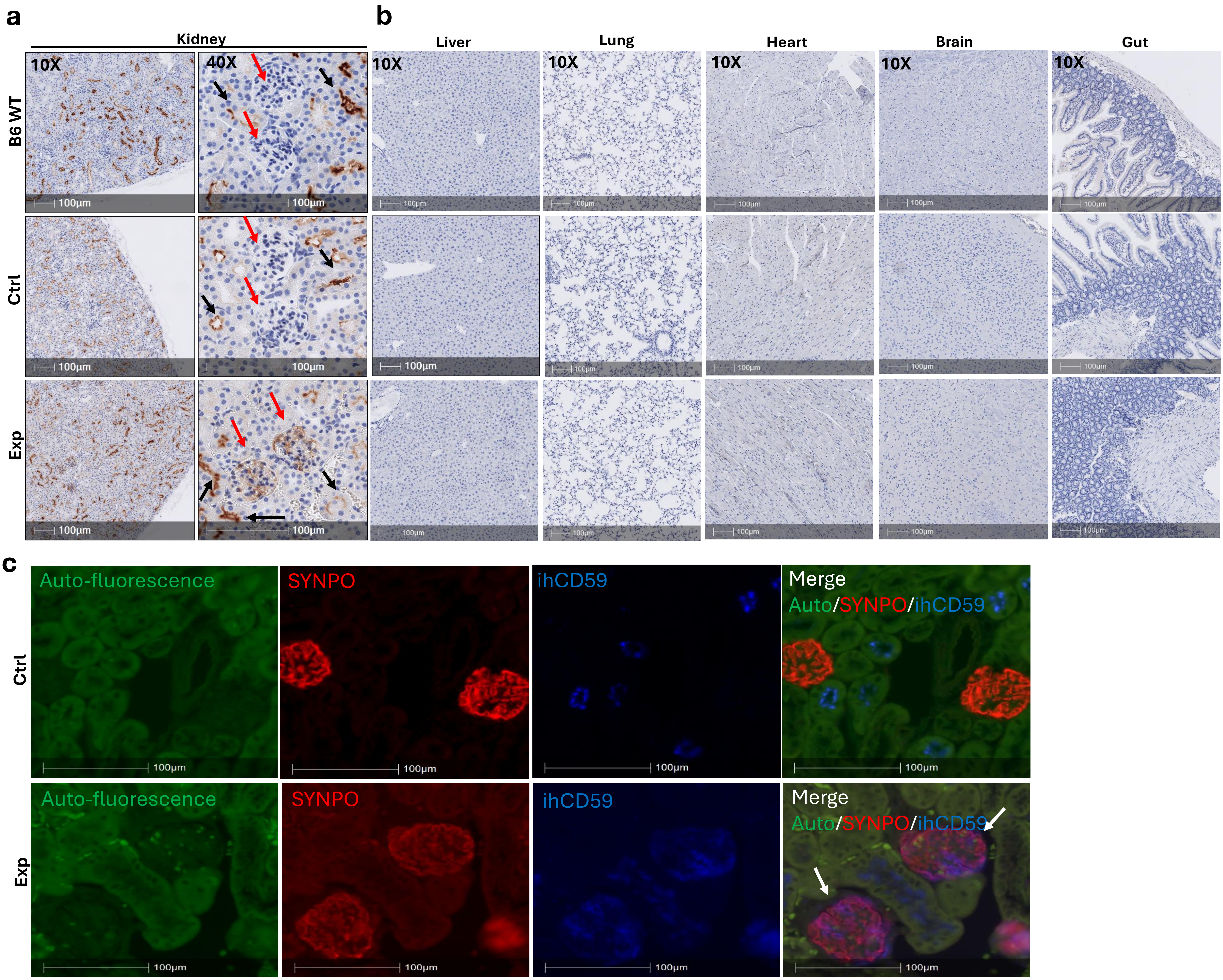


**Supplementary Fig. 1. Specific hCD59 expression on the podocytes of *ihCD59*$^{+/-}$*/Nphs2Cre*$^{+/-}$: (a-b)** Kidneys and major organs (liver, lungs, heart, brain, and intestine) were collected from B6 wild-type (WT, n=1), littermate control (Ctrl) *ihCD59*$^{+/-}$*/Nphs2Cre*$^{-/-}$ (n=1)) and experimental (Exp) *ihCD59*$^{+/-}$*/Nphs2Cre*$^{+/-}$ mice (Exp, n=1). Immunohistochemical (IHC) analysis for hCD59 demonstrated strong and selective expression in the glomeruli (red arrows) of Exp but not Ctrl and B6 mice. Non-specific staining (black arrows) was noted in the brush border of tubule cells across three all mouse genotypes. **(c)** Immunofluorescence (IF) staining confirmed co-localization of hCD59 with the podocyte marker synaptopodin (SYNPO) (white arrow) in glomeruli of Exp (n**=3)** but not Ctrl mice (n=3).

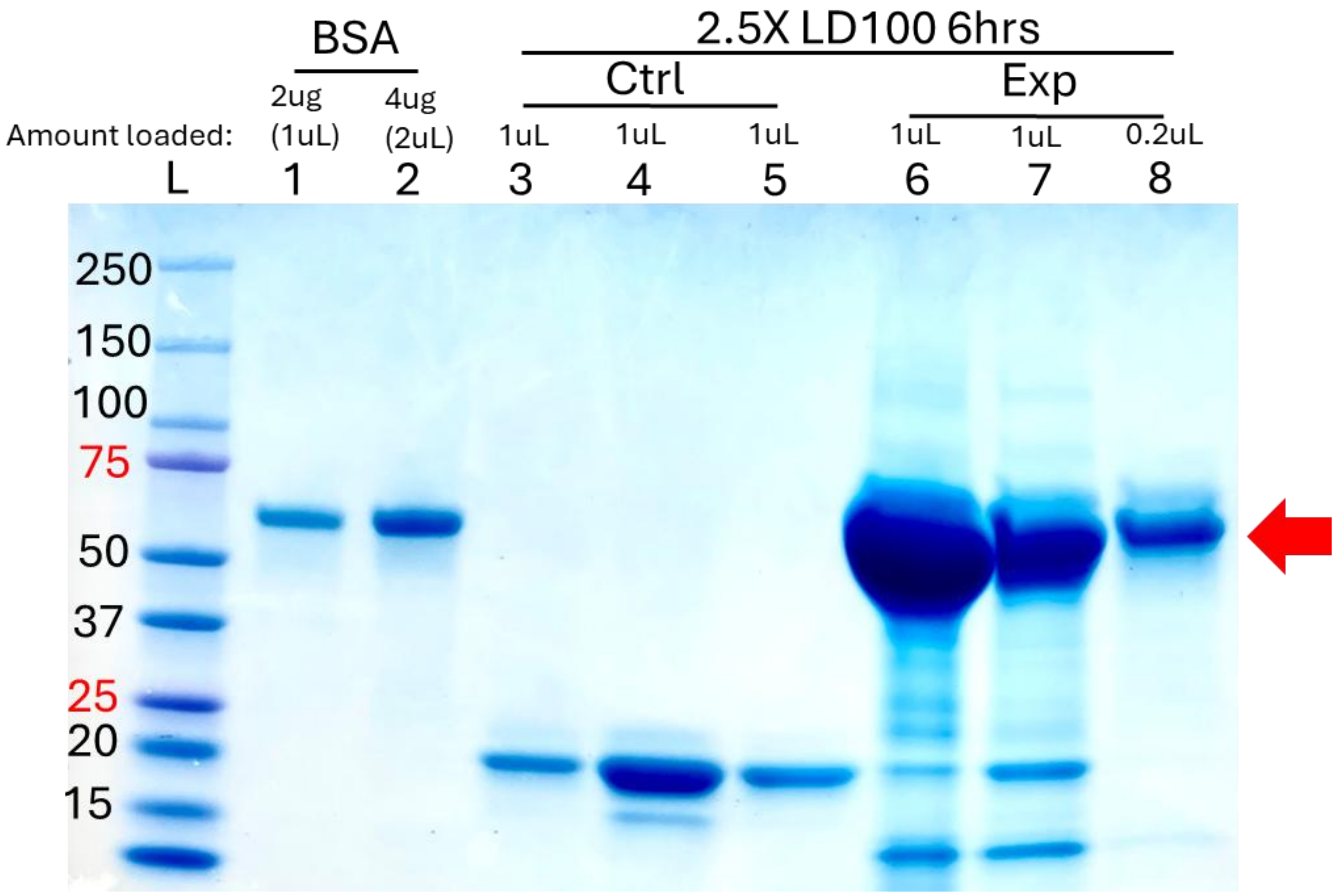


**Supplemental Fig. 2. Uncropped SDS-PAGE image for Fig 1c.**

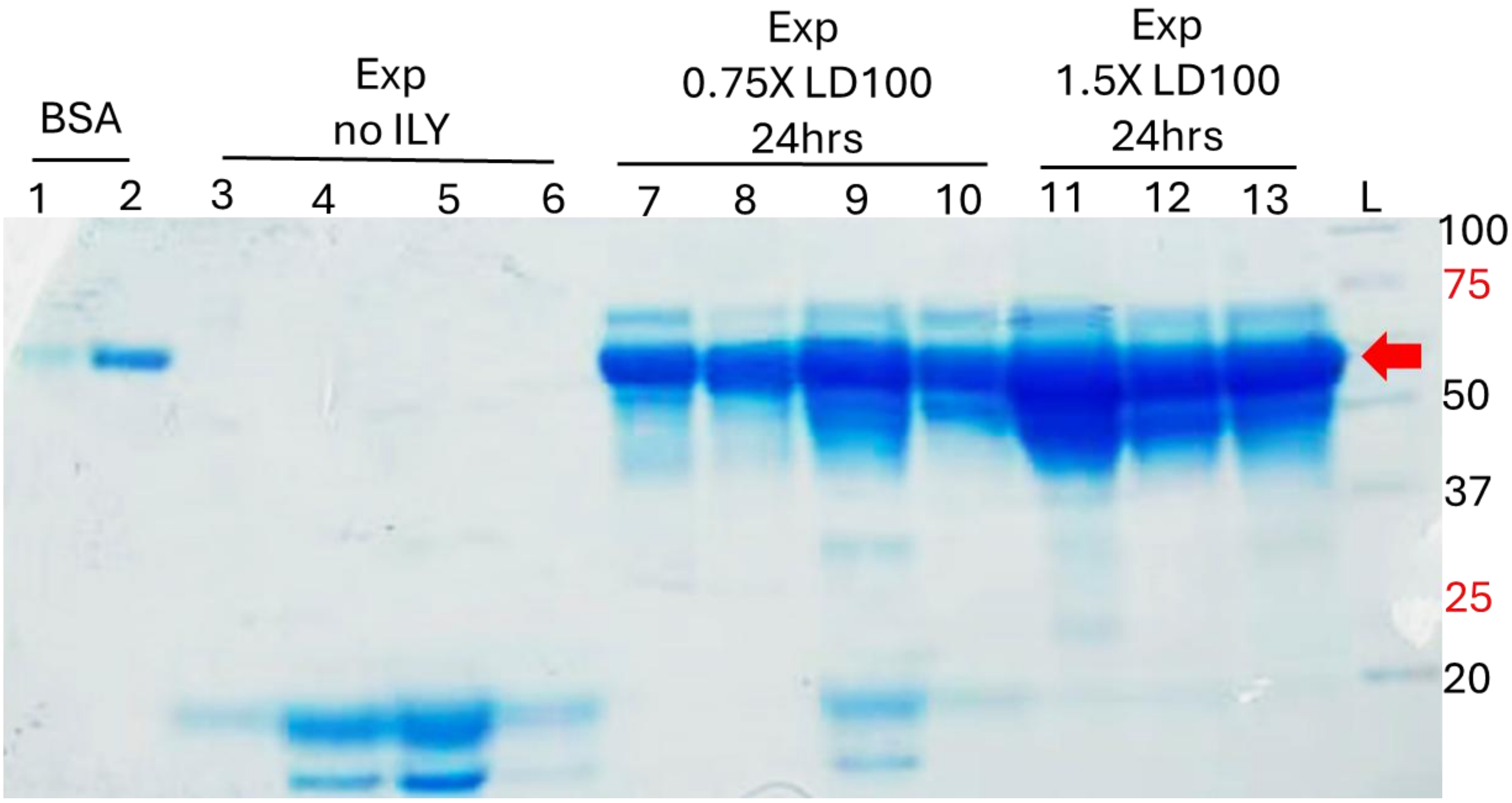


**Supplemental Fig. 3. Uncropped SDS-PAGE image for Fig 3e.**

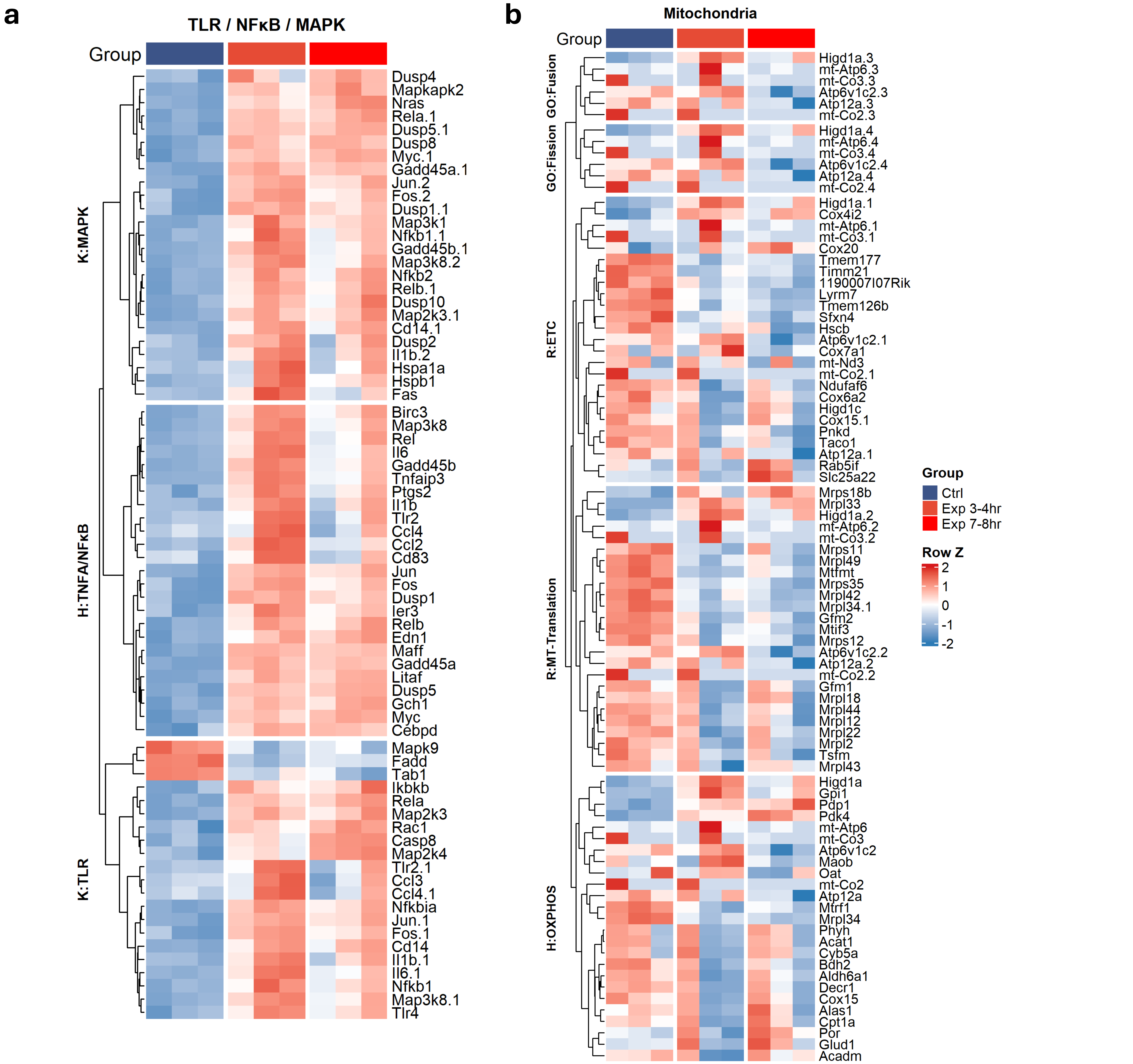


**Supplementary Fig. 4. TLR/NFκB/MAPK signaling and mitochondrial gene expression in the podocyte-ablated mice.(a)** Heatmap showing row-normalized expression (Z-score) of genes within TLR, NFκB, and MAPK-related pathways (n=3 per group). Genes are grouped by pathway annotation: KEGG MAPK signaling (K: MAPK), HALLMARK TNFα/NFκB signaling (H: TNFA/NFκB), and KEGG TLR signaling (K: TLR). Hierarchical clustering was performed within each pathway group. (**b**) Heatmap showing row-normalized expression (Z-score) of mitochondrial genes (n=3 per group). Genes are grouped by pathway annotation: GO Mitochondrial Fusion (GO: Fusion), GO Mitochondrial Fission (GO: Fission), REACTOME Electron Transport Chain (R: ETC), REACTOME Mitochondrial Translation (R: MT-Translation), and HALLMARK Oxidative Phosphorylation (H: OXPHOS). Hierarchical clustering was performed within each pathway group. Color scale represents row Z-score ranging from −2 (blue, downregulated) to +2 (red, upregulated).

Ctrl: Kidneys of littermate control (Ctrl) *ihCD59*$^{+/-}$*/Nphs2Cre*$^{-/-}$ collected at 3-4 hours (n=2) and 7-8 hr (n=1) post ILY injection (1.5XLD100).

Exp 3-4hr: Experimental (*ihCD59*$^{+/-}$*/Nphs2Cre*$^{+/-}$) mice at 3-4 hr post ILY injection (1.5 LD100) (n=3)

Exp 7-8 hr: Experimental (*ihCD59*$^{+/-}$*/Nphs2Cre*$^{+/-}$) mice at 7-8 hr post ILY injection (1.5 LD100, n=3).

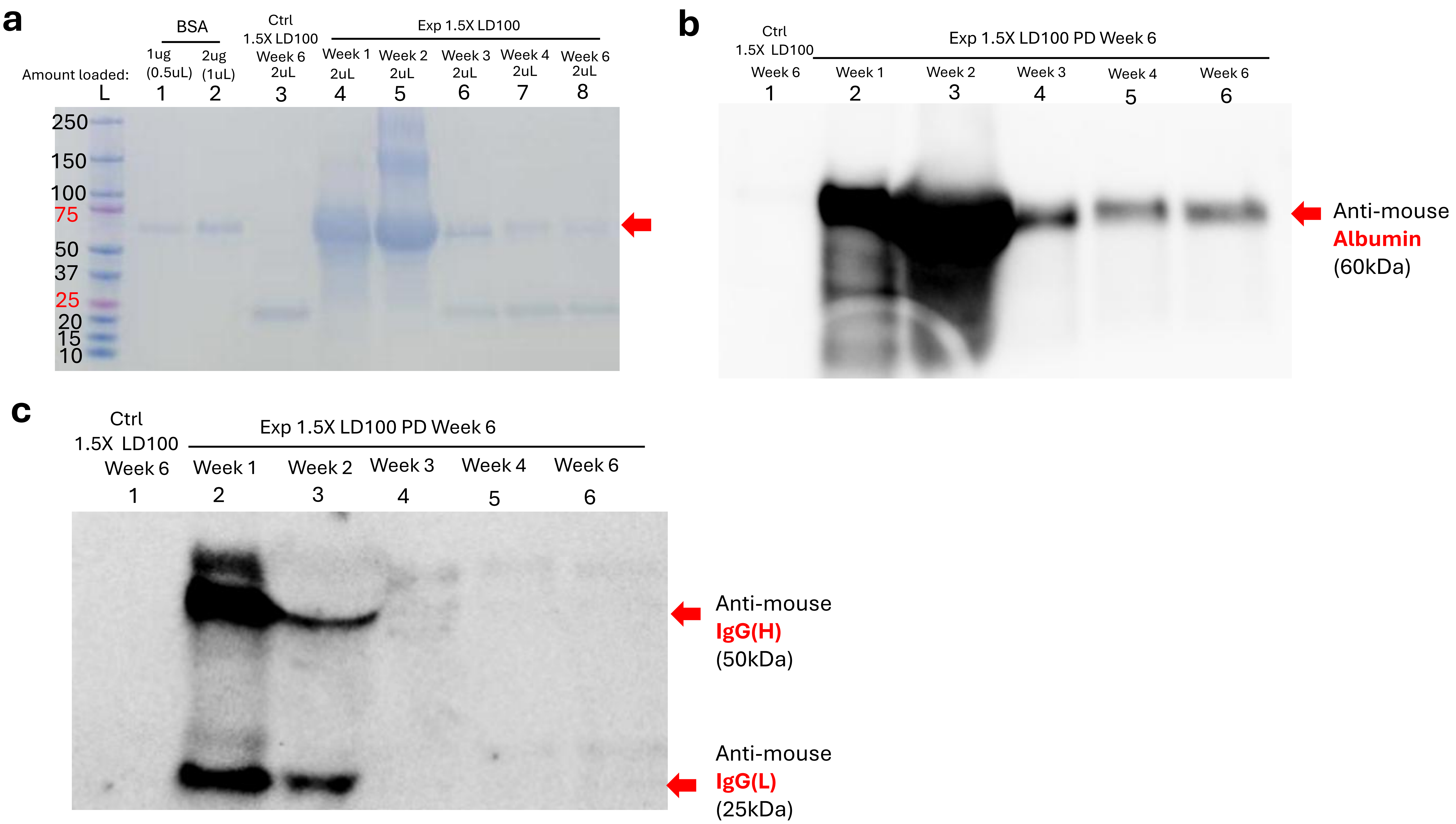


**Supplementary Fig. 5. Uncropped SDS-PAGE image for Fig 6.** (a) Uncropped SDS-PAGE image for Fig. 6a. (b) Uncropped Western Blot image for Fig. 6c. (c) Uncropped Western Blot image for Fig. 6f.

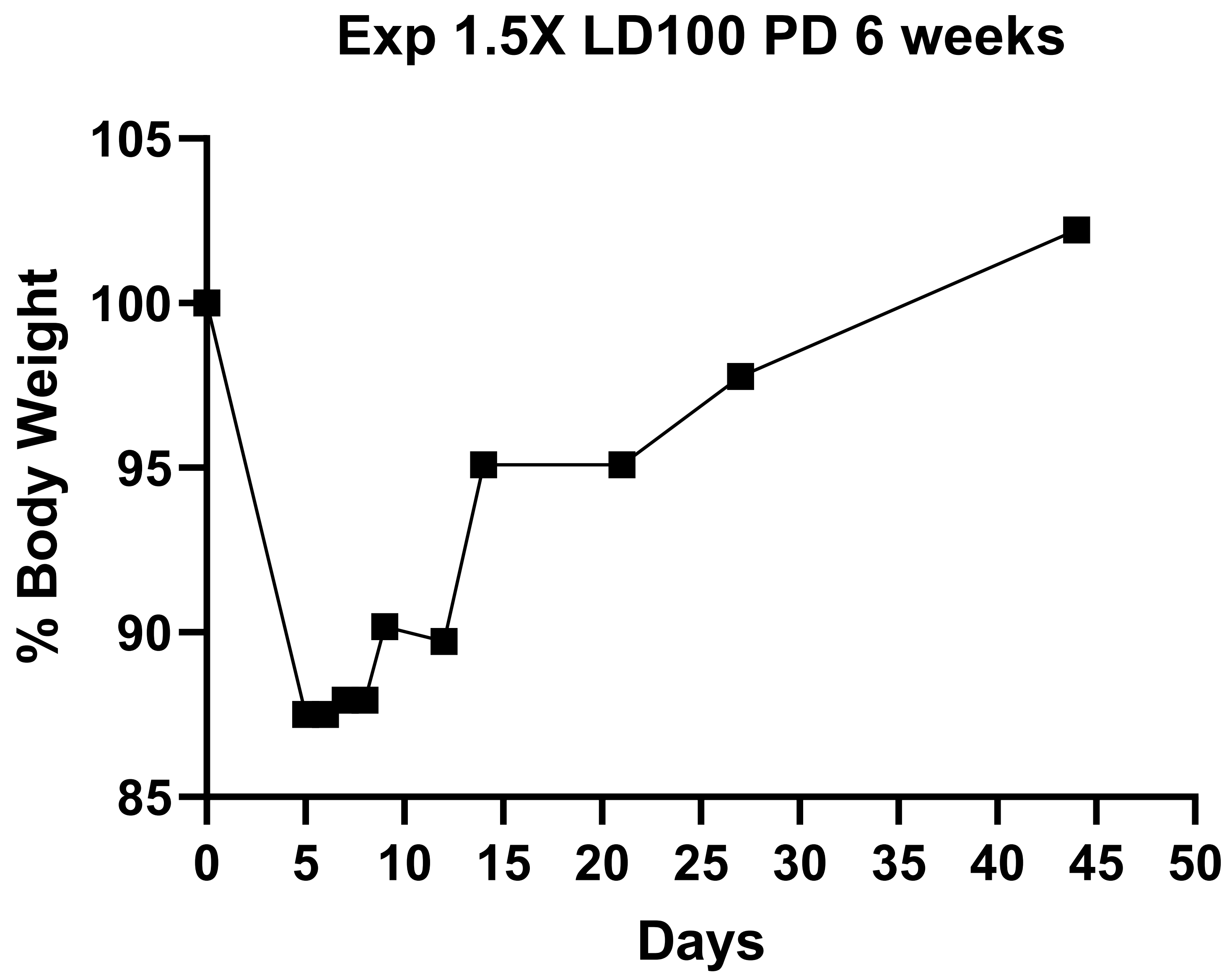


**Supplementary Fig 6. Weight changes in 6 weeks of 1.5X LD100 ILY injected mice + PD treatment.** The percentage of BW change is calculated as the body weight at the given day divided by baseline body weight X 100%.

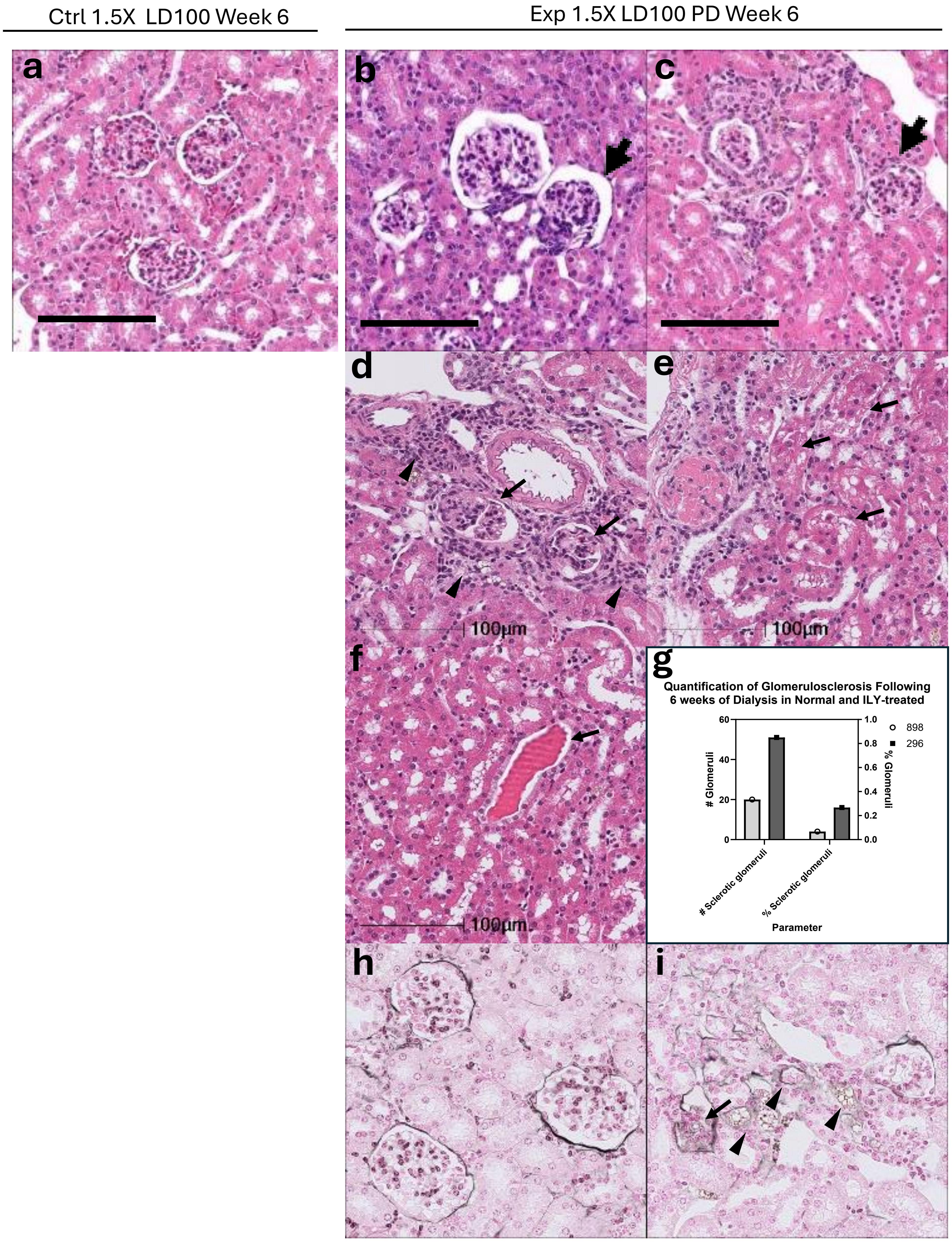


**Supplementary Figure 7: Renal pathology in podocyte-ablated mice after 6 weeks of peritoneal dialysis. (a)** The control mouse has normal glomerular histomorphology. **(b)** Representative images of experimental mice (n=2) with a lethal dose (1.5X LD100), with PD fluid exchanges at 6 weeks, show a sclerotic glomerulus (arrow). **(c)** Multifocal regions of stromal collapse and interstitial fibrosis with some glomeruli exhibit normal morphology (arrow). (**d**) There is multifocal sclerosis of glomeruli (arrows) and regions of tubular loss and replacement by fibrosis (arrowheads). (**e**) Scattered regions of ongoing proximal tubular degeneration (arrows). (**f**) Renal tubules are occasionally dilated by proteinaceous fluid. (**g**) Comparison of the number of sclerotic glomeruli in ILY-, PD-treated (296), and untreated mice (898). **h&i**) PAMS staining demonstrating normal glomerular basement membrane thickness in unablated (**e**) mouse and thickened basement membranes in sclerotic glomeruli (arrow) and in regions of stromal collapse (**f**, arrowheads) in an ILY-, PD-treated mouse. H&E stain. Bar =100 um

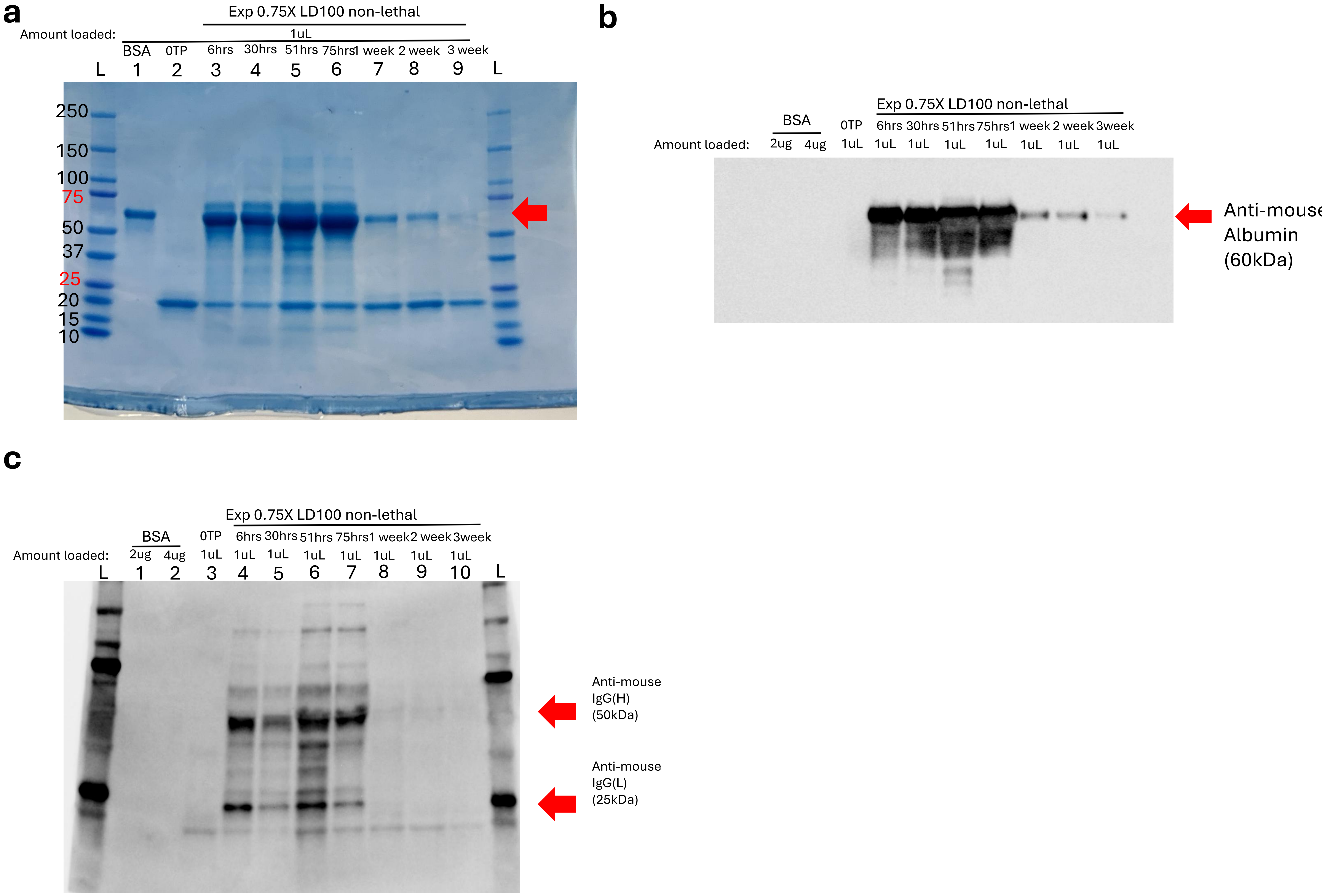


**Supplementary Fig.8.** (**a-b**) Visualization of albuminuria in the experimental mouse received non-lethal dose ILY (0.75X LD100) in week 1-3 by SDS-PAGE **(a)** and western blot (**b**). (**c**) Decreasing infiltrated immunoglobulin (IgG) levels visualized by western blot in the experimental mouse received non-lethal dose ILY (0.75X LD100).

### Supplemental Tables

**Supplementary Table 1: Daily Animal Health Assessment (AHA) Scoring Criteria.**

**Daily Animal Health Assessments Scoring**

| SCORE | DESCRIPTION |
|---|---|
| **1** | • Bright<br>• Alert<br>• responsive |
| **2** | • +/-piloerection<br>• Responsive<br>• ambulatory |
| **3**<br>*If any of these criteria are observed, twice daily monitoring will begin* | • Hunched<br>• decreased activity<br>• responsive to touch<br>• eyes partially closed<br>• unilateral or bilateral ocular discharge<br>• decreased skin turgor |
| **4**<br>*If any three of these criteria are observed, the animal will be euthanized promptly* | • Lethargic<br>• delayed response to touch<br>• delayed righting reflex<br>• opaque eye discharge<br>• cold to the touch<br>• pale pinnae or mucus membranes |
| **5**<br>*If any of these criteria are observed, the animal will be euthanized promptly* | • non-ambulatory<br>• Nonresponsive<br>• no righting reflex<br>• decreased body temperature |

**Supplemental table 2. Bulk RNA-seq mice information**

| # | Eartag | sex | genotype | | group | DOB | Harvest Date | treatment |
|---|---|---|---|---|---|---|---|---|
| | | | Nphs2-Cre | ihCD59 | | | | |
| 1 | 682 | F | -/- | +/- | Control | 6/18/2025 | 09/26/2025 | 3-4hr ILY 1.5X LD100 |
| 2 | 668 | M | -/- | +/- | Control | 6/18/2025 | 09/26/2025 | 3-4hr ILY 1.5X LD100 |
| 3 | 680 | F | +/- | +/- | Exp | 6/18/2025 | 09/26/2025 | 3-4hr ILY 1.5X LD100 |
| 4 | 683 | F | +/- | +/- | Exp | 6/18/2025 | 09/26/2025 | 3-4hr ILY 1.5X LD100 |
| 5 | 669 | M | +/- | +/- | Exp | 6/18/2025 | 09/26/2025 | 3-4hr ILY 1.5X LD100 |
| 6 | 662 | M | -/- | +/- | Control | 6/21/2025 | 09/26/2025 | 7-8hr ILY 1.5X LD100 |
| 7 | 659 | M | +/- | +/- | Exp | 6/21/2025 | 09/26/2025 | 7-8hr ILY 1.5X LD100 |
| 8 | 660 | M | +/- | +/- | Exp | 6/21/2025 | 09/26/2025 | 7-8hr ILY 1.5X LD100 |
| 9 | 665 | F | +/- | +/- | Exp | 6/21/2025 | 09/26/2025 | 7-8hr ILY 1.5X LD100 |

**Supplementary Table 3: Gene Ontology Biology Process showing enriched terms for downregulated DEGs in Exp at 3-4hr post ILY injection vs Control.**

| Term | Count | Genes | Fold Enrichment | PValue | FDR |
|---|---|---|---|---|---|
| GO:0006355~regulation of DNA-templated transcription | 20 | ZFP964, ZFP974, CIITA, L3MBTL3, ZFP961, ZFP213, ZBTB24, PLAG1, ZFP606, ACVR2B, ZFP809, DBP, ZFP85, SUFU, ZFP120, ZFP563, ZFP101, ZFP280B, ZFP551, GM45871 | 2.953802106 | 5.01851E-05 | 0.035179722 |
| GO:0006357~regulation of transcription by RNA polymerase II | 24 | ZFP964, ZFP974, ZFP961, ZFP213, HNF1B, ZFP606, GTF2H2, ZBTB5, AI854703, ZFP809, ZBTB41, GLIS2, DBP, ZKSCAN7, ZFP85, TRPS1, ZFP120, SP5, RCOR3, ZFP563, ZFP750, ZFP101, ZFP551, GM45871 | 2.342709586 | 0.000232423 | 0.081464314 |
| GO:0021532~neural tube patterning | 3 | TBC1D32, IFT140, DZIP1L | 57.43298969 | 0.001145486 | 0.267661918 |
| GO:0007368~determination of left/right symmetry | 5 | ODAD3, TBC1D32, IFT140, SUFU, ACVR2B | 9.009096422 | 0.002318844 | 0.406377385 |
| GO:1905146~lysosomal protein catabolic process | 3 | AP5Z1, VPS13A, MGAT3 | 30.63092784 | 0.004168928 | 0.584483745 |
| GO:0001822~kidney development | 6 | INVS, PTCD2, TBC1D32, HNF1B, ACVR2B, GLIS2 | 4.994173017 | 0.007169538 | 0.837641009 |
| GO:1905349~ciliary transition zone assembly | 2 | DZIP1L, CCDC66 | 102.1030928 | 0.019361415 | 1 |
| GO:0031016~pancreas development | 3 | INVS, HNF1B, ACVR2B | 13.51364463 | 0.02054764 | 1 |
| GO:0061512~protein localization to cilium | 3 | TBC1D32, IFT140, DZIP1L | 11.20643701 | 0.02916047 | 1 |
| GO:0007030~Golgi organization | 4 | UBXN2B, AP5Z1, HIKESHI, CIT | 5.672394044 | 0.033524829 | 1 |
| GO:0007224~smoothened signaling pathway | 4 | TBC1D32, SUFU, DZIP1L, CFAP410 | 5.620353731 | 0.034313753 | 1 |
| GO:0006914~autophagy | 5 | 1600014C10RIK, TRIM17, AP5Z1, VPS13A, PIK3C3 | 4.030385241 | 0.035770257 | 1 |
| GO:0009791~post-embryonic development | 4 | INVS, GRCC10, SZT2, ACVR2B | 5.23605604 | 0.040968431 | 1 |
| GO:0006281~DNA repair | 6 | POLQ, RFC2, MSH3, CHRNA4, APEX2, POLI | 3.157827612 | 0.04178192 | 1 |
| GO:0021540~corpus callosum morphogenesis | 2 | GRCC10, SZT2 | 38.28865979 | 0.050805177 | 1 |
| GO:1905515~non-motile cilium assembly | 3 | BBS10, TBC1D32, IFT140 | 7.179123711 | 0.065117611 | 1 |
| GO:0050890~cognition | 3 | GRCC10, CHRNA4, MGAT3 | 7.179123711 | 0.065117611 | 1 |
| GO:0060271~cilium assembly | 5 | TBC1D32, IFT140, DZIP1L, CCDC66, CFAP410 | 3.300746534 | 0.065169375 | 1 |
| GO:0060287~epithelial cilium movement involved in determination of left/right asymmetry | 2 | INVS, ODAD3 | 27.84629803 | 0.06918785 | 1 |
| GO:0045869~negative regulation of single stranded viral RNA replication via double stranded DNA intermediate | 2 | TASOR, ZFP809 | 27.84629803 | 0.06918785 | 1 |
| GO:0006044~N-acetylglucosamine metabolic process | 2 | GNPDA2, MGAT3 | 19.1443299 | 0.099041883 | 1 |

#### Supplementary Table 4: Summary of Human Glomerulus Dysfunction-related Disease public cohorts analyzed in Fig. 6.

| Accession number | Pathology | Summary |
|---|---|---|
| GSE141295 | Immunoglobulin A nephropathy patients | We collected glomeruli from biopsy specimens from IgA nephropathy patients with relatively preserved kidney function (eGFR ≥ 60 mL/min/1.73 m2 and urine protein-to-creatinine ratio < 3 g/g) and from normal kidney cortices by hand microdissection and performed RNA-seq. The transcriptomic profiling of IgA nephropathy glomerulus provide insights for intraglomerular pathophysiology of IgAN before reaching profound kidney dysfunction. |
| GSE142025 | Diabetic Nephropathy | Comparison of Kidney Transcriptomic Profiles of Early and Advanced Diabetic Nephropathy Reveals Potential New Mechanisms for Disease Progression. Patients with early DN had an eGFR >90 mL/min/1.73 m2 and microalbuminuria (UACR <300 mg/g). Patients with advanced DN had either an eGFR <90 mL/min/1.73 m2 or UACR >300 mg/g. To minimize the tissue processing (i.e., without any dissections or digestions) that may change the gene expression profiles (15), RNA-seq was performed using the whole-kidney biopsy samples, which contain primary kidney cortices. |
| GSE175759 | 43 IgA nephropathy, 3 diabetes mellitus nephropathy, 3 focal segmental glomerulosclerosis, 3 lupus nephritis, 4 membranous nephropathy, and 9 minimal change disease biopsy cores and 22 nephrectomy controls | We profiled manually microdissected tubulointerstitial tissue from 43 IgA nephropathy, 3 diabetes mellitus nephropathy, 3 focal segmental glomerulosclerosis, 3 lupus nephritis, 4 membranous nephropathy, and 9 minimal change disease biopsy cores and 22 nephrectomy controls by RNA sequencing. The 3 outliers which were not included in our main analysis were also uploaded in this database. Patients with profound kidney dysfunction and those with an estimated glomerular filtration rate (eGFR) < 30 mL/min/1.73 m2 were not considered for this study. To establish the control group, normal cortical tissues were collected from the non-cancer-affected cortex of 22 renal carcinoma patients who exhibited no evidence of CKD (eGFR < 60 mL/min/1.73 m2 or detected dipstick albuminuria). These nephrectomy control group samples were also stored in RNAlater immediately after surgical excision via the same protocol used for the biopsy core tissues. The frozen cortical tissues from the nephrectomy controls were initially cut into the size of a biopsy core, and all collected kidney tissues were manually microdissected under a stereomicroscope (Olympus, Japan) to remove glomeruli. After thorough removal of glomeruli, total RNA was extracted from the remaining tubulointerstitial tissues using an RNeasy Micro Kit (Qiagen, Hamburg, Germany). |

#### Supplementary Table 5: TLR genes pan-disease transcriptomic changes.

| Gene | Group | Log2FC | P value | p.adj |
|---|---|---|---|---|
| | Exp 3-4hr_whole_kidney | 1.52261046 | 0.0040402 | 0.04748356 |
| | Exp 7-8hr_whole_kidney | 0.92826276 | 0.10146881 | 0.25379422 |
| | DN - Adv_whole_kidney | 0.70971932 | 5.7714E-06 | 3.4107E-05 |
| | IgAN - Glom | 1.45069531 | 0.00150326 | 0.00835348 |
| | IgAN - TI | 0.47496259 | 0.00357766 | 0.02562322 |
| | FSGS - TI | 1.4841272 | 0.00011263 | 0.00164522 |
| | LN - TI | 1.7889272 | 3.1739E-06 | 0.00024328 |
| | MN - TI | 1.05635361 | 0.00188692 | 0.04562147 |
| **TLR2** | MCD - TI | 0.73267844 | 0.00313222 | 0.05257494 |
| | Exp 3-4hr_whole_kidney | 1.55361686 | 0.00030413 | 0.00892143 |
| | Exp 7-8hr_whole_kidney | 1.37276985 | 0.00282557 | 0.02214834 |
| | DN - Adv_whole_kidney | 0.5102827 | 4.164E-06 | 2.5541E-05 |
| | IgAN - Glom | 0.38463107 | 0.1217056 | 0.22863175 |
| | IgAN - TI | 0.33488587 | 0.02415035 | 0.09379977 |
| | FSGS - TI | 1.28135946 | 0.00026808 | 0.00310751 |
| | LN - TI | 1.18092974 | 0.00078378 | 0.01667149 |
| | MN - TI | 0.73107802 | 0.01866889 | 0.15437059 |
| **TLR4** | MCD - TI | 0.04958863 | 0.82681911 | 0.9362462 |
| | Exp 3-4hr_whole_kidney | 2.2128265 | 0.00013063 | 0.00507247 |
| | Exp 7-8hr_whole_kidney | 2.22212034 | 0.00029904 | 0.00452622 |
| | DN - Adv_whole_kidney | 1.95725595 | 2.8049E-11 | 6.4609E-10 |
| | IgAN - Glom | 1.3969705 | 0.06595984 | 0.14435816 |
| | IgAN - TI | 1.18512394 | 4.6122E-05 | 0.00116819 |
| | FSGS - TI | 1.91953061 | 0.00444896 | 0.02582251 |
| | LN - TI | 3.60933858 | 7.1042E-08 | 1.1674E-05 |
| | MN - TI | 2.33698688 | 8.58E-05 | 0.00763459 |
| **TLR8** | MCD - TI | 0.54942486 | 0.21337235 | 0.53574642 |

#### Supplementary Table 6: Mitochondrial function related genes pan-disease transcriptomic changes.

| gene | dataset_label | log2FoldChange | pvalue | padj |
|---|---|---|---|---|
| | Exp 3-4hr_whole_kidney | 5.79547687 | NA | NA |
| | Exp 7-8hr_whole_kidney | -0.72100348 | NA | NA |
| | DN - Adv_whole_kidney | -3.53457425 | 1.1542E-11 | 2.9367E-10 |
| | IgAN - Glom | -0.36404786 | 0.04703768 | 0.11252718 |
| | IgAN - TI | -0.66730707 | 2.2953E-06 | 0.00011433 |
| | FSGS - TI | -1.11476454 | 0.00088466 | 0.00758282 |
| | LN - TI | -1.55792668 | 3.3745E-06 | 0.00025498 |
| | MN - TI | -1.42499636 | 1.4918E-06 | 0.00054937 |
| MT-ATP6 | MCD - TI | -1.23240332 | 1.0822E-08 | 5.6506E-06 |
| | Exp 3-4hr_whole_kidney | -1.27095471 | NA | NA |
| | Exp 7-8hr_whole_kidney | -23.2588375 | NA | NA |
| | DN - Adv_whole_kidney | -2.56370763 | 2.5934E-15 | 1.71E-13 |
| | IgAN - Glom | -0.49589736 | 0.01168477 | 0.04007014 |
| | IgAN - TI | -0.74083506 | 1.7528E-07 | 1.4167E-05 |
| | FSGS - TI | -1.41899825 | 2.507E-05 | 0.00053965 |
| | LN - TI | -1.56455023 | 3.3772E-06 | 0.00025498 |
| | MN - TI | -1.54415353 | 2.0753E-07 | 0.00010137 |
| MT-CO2 | MCD - TI | -1.1850616 | 4.3945E-08 | 1.5878E-05 |
| | Exp 3-4hr_whole_kidney | -0.7240668 | NA | NA |
| | Exp 7-8hr_whole_kidney | -8.06194458 | NA | NA |
| | DN - Adv_whole_kidney | -1.83910772 | 4.1581E-12 | 1.1979E-10 |
| | IgAN - Glom | -0.77925012 | 0.00169032 | 0.00914005 |
| | IgAN - TI | -0.73080029 | 3.6332E-07 | 2.5193E-05 |
| | FSGS - TI | -1.63518638 | 1.6333E-06 | 7.8599E-05 |
| | LN - TI | -1.74857065 | 2.9499E-07 | 3.8428E-05 |
| | MN - TI | -1.56719916 | 1.9646E-07 | 9.7968E-05 |
| MT-CO3 | MCD - TI | -1.18062946 | 7.2812E-08 | 2.4817E-05 |
| | Exp 3-4hr_whole_kidney | 0.32044828 | 0.20816861 | 0.43351251 |
| | Exp 7-8hr_whole_kidney | 0.00196592 | 0.99423711 | 0.99741092 |
| | DN - Adv_whole_kidney | -2.58312374 | 6.9028E-15 | 4.0715E-13 |
| | IgAN - Glom | -0.5331663 | 0.02016573 | 0.06025044 |
| | IgAN - TI | -0.61366274 | 2.6771E-05 | 0.00077092 |
| | FSGS - TI | -1.30693161 | 0.00016538 | 0.00217613 |
| | LN - TI | -1.57038686 | 6.0092E-06 | 0.00041503 |
| | MN - TI | -1.43036284 | 3.0442E-06 | 0.00094888 |
| MT-CYB | MCD - TI | -1.36273865 | 1.0019E-09 | 8.7549E-07 |
| | Exp 3-4hr_whole_kidney | -0.31911864 | 0.84617115 | NA |
| | Exp 7-8hr_whole_kidney | -0.49967138 | 0.77908387 | NA |
| | DN - Adv_whole_kidney | -2.95051436 | 4.8985E-23 | 2.4537E-20 |
| | IgAN - Glom | -0.23381592 | 0.40643036 | 0.54545504 |
| | IgAN - TI | -0.44273147 | 0.00182879 | 0.01601426 |
| | FSGS - TI | -1.10317728 | 0.0010722 | 0.00877644 |
| | LN - TI | -1.47724004 | 1.1873E-05 | 0.0006894 |
| | MN - TI | -1.44359345 | 1.2577E-06 | 0.00047784 |
| MT-ND3 | MCD - TI | -1.02062533 | 2.515E-06 | 0.00037633 |
| | Exp 3-4hr_whole_kidney | 0.10134013 | 0.65915465 | 0.80932342 |
| | Exp 7-8hr_whole_kidney | -0.16327775 | 0.50620729 | 0.69051659 |
| | DN - Adv_whole_kidney | -3.11591813 | 1.4482E-10 | 2.7821E-09 |
| | IgAN - Glom | -0.36943241 | 0.04738707 | 0.11317875 |
| | IgAN - TI | -0.60567501 | 8.5586E-06 | 0.00032588 |
| | FSGS - TI | -1.11787065 | 0.00054064 | 0.00531756 |
| | LN - TI | -1.34822637 | 3.01E-05 | 0.00139369 |
| | MN - TI | -1.31038953 | 4.3887E-06 | 0.00118032 |
| MT-ND4 | MCD - TI | -1.03550063 | 6.1996E-07 | 0.00013125 |

**Supplementary Table 7: Consistently Upregulated TLR related pathways across podocyte ablated mice and human glomerulus dysfunction related disease cohorts via GSEA analysis**

| Description | dataset_label | NES | pvalue | qvalue | setSize | sig_label |
|---|---|---|---|---|---|---|
| GOBP_TOLL_LIKE_RECEPTOR_SIGNALING_PATHWAY | Exp_3_4_hr_Whole_kidney | 1.89399306 | 9.4447E-06 | 8.2751E-05 | 79 | *** |
| | DN_Whole_kidney | 1.62597531 | 0.00054188 | 0.00736559 | 84 | ** |
| | LN_TI | 2.5158784 | 1E-10 | 3.4331E-09 | 82 | *** |
| | MN_TI | 1.95300165 | 1.8534E-05 | 0.00048653 | 82 | *** |
| | MCD_TI | 1.63419481 | 0.00136447 | 0.02675577 | 82 | * |
| GOBP_MYD88_DEPENDENT_TOLL_LIKE_RECEPTOR_SIGNALING_PATHWAY | Exp_3_4_hr_Whole_kidney | 1.63194151 | 0.01000132 | 0.02397471 | 21 | * |
| | IgAN_Glom. | 2.42540129 | 2.372E-05 | 0.00384759 | 20 | ** |
| | FSGS_TI | 1.91479577 | 0.00076297 | 0.00523396 | 31 | ** |
| | LN_TI | 2.15417867 | 1.2829E-05 | 0.00015707 | 21 | *** |
| | MN_TI | 1.98329539 | 0.00037142 | 0.00612998 | 21 | ** |
| | MCD_TI | 1.85830403 | 0.00216922 | 0.03717008 | 21 | * |
| GOBP_ENDOLYSOSOMAL_TOLL_LIKE_RECEPTOR_SIGNALING_PATHWAY | Exp_3_4_hr_Whole_kidney | 1.70938545 | 0.00088018 | 0.00357948 | 55 | ** |
| | IgAN_TI | 1.92716163 | 8.4794E-05 | 0.00369545 | 52 | ** |
| | LN_TI | 2.33078072 | 6.2818E-09 | 1.5801E-07 | 52 | *** |
| | MN_TI | 2.21896756 | 4.0521E-07 | 1.6963E-05 | 52 | *** |
| | MCD_TI | 1.92978481 | 0.00020009 | 0.00677384 | 52 | ** |

**Supplementary Table 8: Consistently downregulated mitochondria function-related pathways across podocyte-ablated mice and human glomerulus dysfunction-related disease cohorts via GSEA analysis.**

| Description | dataset_label | NES | pvalue | qvalue | setSize | sig_label |
|---|---|---|---|---|---|---|
| HP_MITOCHONDRIAL_INHERITANCE | Exp_3_4_hr_Whole_kidney | -1.94944 | 0.001271 | 0.004788 | 15 | ** |
| | Exp_7_8_hr_Whole_kidney | -2.1531 | 9.39E-08 | 6.39E-05 | 15 | *** |
| | DN_Whole_kidney | -2.9082 | 1E-10 | 1.13E-08 | 28 | *** |
| | FSGS_TI | -1.74991 | 0.003495 | 0.018745 | 28 | * |
| | LN_TI | -2.00112 | 0.000161 | 0.001476 | 28 | ** |
| | MN_TI | -2.21944 | 1.6E-06 | 5.62E-05 | 28 | *** |
| | MCD_TI | -2.03152 | 1.71E-05 | 0.001063 | 28 | ** |
| HP_MITOCHONDRIAL_RESPIRATORY_CHAIN_DEFECTS | Exp_7_8_hr_Whole_kidney | -1.97043 | 1.63E-05 | 0.003699 | 18 | ** |
| | DN_Whole_kidney | -2.58605 | 6.35E-10 | 6.13E-08 | 19 | *** |
| | LN_TI | -1.87867 | 0.001564 | 0.010313 | 19 | * |
| | MN_TI | -2.04007 | 0.000155 | 0.002974 | 19 | ** |
| | MCD_TI | -1.87621 | 0.000477 | 0.013014 | 19 | * |
| HP_ABNORMAL_MITOCHONDRIA_IN_MUSCLE_TISSUE | Exp_3_4_hr_Whole_kidney | -1.995 | 0.00035 | 0.001687 | 39 | ** |
| | Exp_7_8_hr_Whole_kidney | -2.48272 | 1E-10 | 2.31E-07 | 39 | *** |
| | DN_Whole_kidney | -2.38431 | 1.27E-07 | 6.2E-06 | 47 | *** |
| | IgAN_TI | -1.70769 | 0.002573 | 0.042025 | 47 | * |
| | FSGS_TI | -2.00159 | 0.000235 | 0.001929 | 47 | ** |
| | LN_TI | -1.84827 | 0.000403 | 0.00325 | 47 | ** |
| | MN_TI | -2.17145 | 1.01E-05 | 0.000291 | 47 | *** |
| | MCD_TI | -1.92553 | 6.53E-05 | 0.002897 | 47 | ** |
| HP_MITOCHONDRIAL_MYOPATHY | Exp_3_4_hr_Whole_kidney | -1.68786 | 0.003783 | 0.011427 | 54 | * |
| | Exp_7_8_hr_Whole_kidney | -2.4474 | 1E-10 | 2.31E-07 | 54 | *** |
| | DN_Whole_kidney | -2.7838 | 1E-10 | 1.13E-08 | 63 | *** |
| | IgAN_TI | -1.79041 | 0.000453 | 0.012675 | 63 | * |
| | FSGS_TI | -1.82334 | 0.000589 | 0.004231 | 63 | ** |
| | LN_TI | -1.73544 | 0.001689 | 0.011008 | 63 | * |
| | MN_TI | -2.2424 | 1.12E-07 | 5.54E-06 | 63 | *** |
| | MCD_TI | -1.81362 | 0.000244 | 0.007985 | 63 | ** |
| GOBP_MITOCHONDRIAL_ELECTRON_TRANSPORT_CYTOCHROME_C_TO_OXYGEN | Exp_7_8_hr_Whole_kidney | -2.12516 | 1.74E-05 | 0.003807 | 19 | ** |
| | DN_Whole_kidney | -2.04783 | 0.00057 | 0.007669 | 20 | ** |
| | IgAN_TI | -2.1283 | 1.14E-05 | 0.000707 | 20 | *** |
| | FSGS_TI | -2.20369 | 4.02E-05 | 0.000406 | 20 | *** |
| | LN_TI | -1.77719 | 0.007283 | 0.036713 | 20 | * |
| | MN_TI | -1.95983 | 0.000335 | 0.005587 | 20 | ** |
| HP_MUSCLE_ABNORMALITY_RELATED_TO_MITOCHONDRIAL_DYSFUNCTION | Exp_7_8_hr_Whole_kidney | -2.30173 | 5.7E-09 | 7.33E-06 | 87 | *** |
| | DN_Whole_kidney | -2.61817 | 1E-10 | 1.13E-08 | 100 | *** |
| | IgAN_TI | -1.80691 | 0.000174 | 0.006318 | 100 | ** |
| | FSGS_TI | -1.71648 | 0.001104 | 0.007147 | 100 | ** |
| | LN_TI | -1.63066 | 0.002655 | 0.015916 | 100 | * |
| | MN_TI | -2.12447 | 7.76E-07 | 2.93E-05 | 100 | *** |
| | MCD_TI | -1.6447 | 0.001195 | 0.024789 | 100 | * |

**Supplementary Table 9: Consistently Upregulated Apoptosis pathways across podocyte-ablated mice and human glomerulus dysfunction-related disease cohorts via GSEA analysis**

| Description | dataset_label | NES | pvalue | qvalue | setSize | sig_label |
|---|---|---|---|---|---|---|
| GOBP_REGULATION_OF_EXTRINSIC_APOPTOTIC_SIGNALING_PATHWAY | Exp_3_4_hr_Whole_kidney | 1.97407 | 4.71E-09 | 1.13E-07 | 157 | *** |
| | IgAN_TI | 1.524647 | 0.00261 | 0.042311 | 167 | * |
| | LN_TI | 1.826985 | 7.53E-06 | 9.85E-05 | 167 | *** |
| | MN_TI | 1.49422 | 0.001601 | 0.019791 | 167 | * |
| | MCD_TI | 1.642373 | 0.000126 | 0.00471 | 167 | ** |
| GOBP_EXTRINSIC_APOPTOTIC_SIGNALING_PATHWAY | Exp_3_4_hr_Whole_kidney | 2.071804 | 1E-10 | 3.81E-09 | 225 | *** |
| | LN_TI | 1.931739 | 1.18E-08 | 2.83E-07 | 239 | *** |
| | MN_TI | 1.431781 | 0.002525 | 0.028912 | 239 | * |
| | MCD_TI | 1.565392 | 9.51E-05 | 0.003852 | 239 | ** |
| GOBP_APOPTOTIC_CELL_CLEARANCE | Exp_3_4_hr_Whole_kidney | 1.672703 | 0.002102 | 0.007155 | 37 | ** |
| | LN_TI | 2.300509 | 4.68E-08 | 1.01E-06 | 44 | *** |
| | MN_TI | 2.087208 | 7.85E-06 | 0.000229 | 44 | *** |
| | MCD_TI | 1.792683 | 0.001601 | 0.03031 | 44 | * |

**Supplementary Table 10: Primary antibodies used for the immunohistochemistry studies.**

| PRIMARY ANTIBODY | SPECIES | COMPANY | CATALOG | DILUTION | SECONDARY ANTIBODY | COLOR DEVELOPMENT |
|---|---|---|---|---|---|---|
| CD59 | Rabbit | Sigma | HPA026494 | 1:200 | OMap anti-Rabbit HRP | Discovery Rhodamine |
| Synaptopodin | Rabbit | ProteinTech | 21064-1-AP | 1:400 | OMap anti-Rabbit HRP | Discovery Cy5 |
| Dapi | | Invitrogen | D1306 | | | |

#### Supplementary Table 1: Daily Animal Health Assessment (AHA) Scoring Criteria.

**Daily Animal Health Assessments Scoring**

| SCORE | DESCRIPTION |
|---|---|
| **1** | • Bright<br>• Alert<br>• responsive |
| **2** | • +/-piloerection<br>• Responsive<br>• ambulatory |
| **3**<br>*If any of these criteria are observed, twice daily monitoring will begin* | • Hunched<br>• decreased activity<br>• responsive to touch<br>• eyes partially closed<br>• unilateral or bilateral ocular discharge<br>• decreased skin turgor |
| **4**<br>*If any three of these criteria are observed, the animal will be euthanized promptly* | • Lethargic<br>• delayed response to touch<br>• delayed righting reflex<br>• opaque eye discharge<br>• cold to the touch<br>• pale pinnae or mucus membranes |
| **5**<br>*If any of these criteria are observed, the animal will be euthanized promptly* | • non-ambulatory<br>• Nonresponsive<br>• no righting reflex<br>• decreased body temperature |

**Supplemental table 2. Bulk RNA-seq mice information**

| # | Eartag | sex | genotype | | group | DOB | Harvest Date | treatment |
|---|---|---|---|---|---|---|---|---|
| | | | Nphs2-Cre | ihCD59 | | | | |
| 1 | 682 | F | -/- | +/- | Control | 6/18/2025 | 09/26/2025 | 3-4hr ILY 1.5X LD100 |
| 2 | 668 | M | -/- | +/- | Control | 6/18/2025 | 09/26/2025 | 3-4hr ILY 1.5X LD100 |
| 3 | 680 | F | +/- | +/- | Exp | 6/18/2025 | 09/26/2025 | 3-4hr ILY 1.5X LD100 |
| 4 | 683 | F | +/- | +/- | Exp | 6/18/2025 | 09/26/2025 | 3-4hr ILY 1.5X LD100 |
| 5 | 669 | M | +/- | +/- | Exp | 6/18/2025 | 09/26/2025 | 3-4hr ILY 1.5X LD100 |
| 6 | 662 | M | -/- | +/- | Control | 6/21/2025 | 09/26/2025 | 7-8hr ILY 1.5X LD100 |
| 7 | 659 | M | +/- | +/- | Exp | 6/21/2025 | 09/26/2025 | 7-8hr ILY 1.5X LD100 |
| 8 | 660 | M | +/- | +/- | Exp | 6/21/2025 | 09/26/2025 | 7-8hr ILY 1.5X LD100 |
| 9 | 665 | F | +/- | +/- | Exp | 6/21/2025 | 09/26/2025 | 7-8hr ILY 1.5X LD100 |

**Supplementary Table 3: Gene Ontology Biology Process showing enriched terms for downregulated DEGs in Exp at 3-4hr post ILY injection vs Control.**

| Term | Count | Genes | Fold Enrichment | PValue | FDR |
|---|---|---|---|---|---|
| GO:0006355~regulation of DNA-templated transcription | 20 | ZFP964, ZFP974, CIITA, L3MBTL3, ZFP961, ZFP213, ZBTB24, PLAG1, ZFP606, ACVR2B, ZFP809, DBP, ZFP85, SUFU, ZFP120, ZFP563, ZFP101, ZFP280B, ZFP551, GM45871 | 2.953802106 | 5.01851E-05 | 0.035179722 |
| GO:0006357~regulation of transcription by RNA polymerase II | 24 | ZFP964, ZFP974, ZFP961, ZFP213, HNF1B, ZFP606, GTF2H2, ZBTB5, AI854703, ZFP809, ZBTB41, GLIS2, DBP, ZKSCAN7, ZFP85, TRPS1, ZFP120, SP5, RCOR3, ZFP563, ZFP750, ZFP101, ZFP551, GM45871 | 2.342709586 | 0.000232423 | 0.081464314 |
| GO:0021532~neural tube patterning | 3 | TBC1D32, IFT140, DZIP1L | 57.43298969 | 0.001145486 | 0.267661918 |
| GO:0007368~determination of left/right symmetry | 5 | ODAD3, TBC1D32, IFT140, SUFU, ACVR2B | 9.009096422 | 0.002318844 | 0.406377385 |
| GO:1905146~lysosomal protein catabolic process | 3 | AP5Z1, VPS13A, MGAT3 | 30.63092784 | 0.004168928 | 0.584483745 |
| GO:0001822~kidney development | 6 | INVS, PTCD2, TBC1D32, HNF1B, ACVR2B, GLIS2 | 4.994173017 | 0.007169538 | 0.837641009 |
| GO:1905349~ciliary transition zone assembly | 2 | DZIP1L, CCDC66 | 102.1030928 | 0.019361415 | 1 |
| GO:0031016~pancreas development | 3 | INVS, HNF1B, ACVR2B | 13.51364463 | 0.02054764 | 1 |
| GO:0061512~protein localization to cilium | 3 | TBC1D32, IFT140, DZIP1L | 11.20643701 | 0.02916047 | 1 |
| GO:0007030~Golgi organization | 4 | UBXN2B, AP5Z1, HIKESHI, CIT | 5.672394044 | 0.033524829 | 1 |
| GO:0007224~smoothened signaling pathway | 4 | TBC1D32, SUFU, DZIP1L, CFAP410 | 5.620353731 | 0.034313753 | 1 |
| GO:0006914~autophagy | 5 | 1600014C10RIK, TRIM17, AP5Z1, VPS13A, PIK3C3 | 4.030385241 | 0.035770257 | 1 |
| GO:0009791~post-embryonic development | 4 | INVS, GRCC10, SZT2, ACVR2B | 5.23605604 | 0.040968431 | 1 |
| GO:0006281~DNA repair | 6 | POLQ, RFC2, MSH3, CHRNA4, APEX2, POLI | 3.157827612 | 0.04178192 | 1 |
| GO:0021540~corpus callosum morphogenesis | 2 | GRCC10, SZT2 | 38.28865979 | 0.050805177 | 1 |
| GO:1905515~non-motile cilium assembly | 3 | BBS10, TBC1D32, IFT140 | 7.179123711 | 0.065117611 | 1 |
| GO:0050890~cognition | 3 | GRCC10, CHRNA4, MGAT3 | 7.179123711 | 0.065117611 | 1 |
| GO:0060271~cilium assembly | 5 | TBC1D32, IFT140, DZIP1L, CCDC66, CFAP410 | 3.300746534 | 0.065169375 | 1 |
| GO:0060287~epithelial cilium movement involved in determination of left/right asymmetry | 2 | INVS, ODAD3 | 27.84629803 | 0.06918785 | 1 |
| GO:0045869~negative regulation of single stranded viral RNA replication via double stranded DNA intermediate | 2 | TASOR, ZFP809 | 27.84629803 | 0.06918785 | 1 |
| GO:0006044~N-acetylglucosamine metabolic process | 2 | GNPDA2, MGAT3 | 19.1443299 | 0.099041883 | 1 |

**Supplementary Table 4: Summary of Human Glomerulus Dysfunction-related Disease public cohorts analyzed in Fig. 6.**

| Accession number | Pathology | Summary |
|---|---|---|
| GSE141295 | Immunoglobulin A nephropathy patients | We collected glomeruli from biopsy specimens from IgA nephropathy patients with relatively preserved kidney function (eGFR ≥ 60 mL/min/1.73 m2 and urine protein-to-creatinine ratio < 3 g/g) and from normal kidney cortices by hand microdissection and performed RNA-seq. The transcriptomic profiling of IgA nephropathy glomerulus provide insights for intraglomerular pathophysiology of IgAN before reaching profound kidney dysfunction. |
| GSE142025 | Diabetic Nephropathy | Comparison of Kidney Transcriptomic Profiles of Early and Advanced Diabetic Nephropathy Reveals Potential New Mechanisms for Disease Progression. Patients with early DN had an eGFR >90 mL/min/1.73 m2 and microalbuminuria (UACR <300 mg/g). Patients with advanced DN had either an eGFR <90 mL/min/1.73 m2 or UACR >300 mg/g. To minimize the tissue processing (i.e., without any dissections or digestions) that may change the gene expression profiles (15), RNA-seq was performed using the whole-kidney biopsy samples, which contain primary kidney cortices. |
| GSE175759 | 43 IgA nephropathy, 3 diabetes mellitus nephropathy, 3 focal segmental glomerulosclerosis, 3 lupus nephritis, 4 membranous nephropathy, and 9 minimal change disease biopsy cores and 22 nephrectomy controls | We profiled manually microdissected tubulointerstitial tissue from 43 IgA nephropathy, 3 diabetes mellitus nephropathy, 3 focal segmental glomerulosclerosis, 3 lupus nephritis, 4 membranous nephropathy, and 9 minimal change disease biopsy cores and 22 nephrectomy controls by RNA sequencing. The 3 outliers which were not included in our main analysis were also uploaded in this database. Patients with profound kidney dysfunction and those with an estimated glomerular filtration rate (eGFR) < 30 mL/min/1.73 m2 were not considered for this study. To establish the control group, normal cortical tissues were collected from the non-cancer-affected cortex of 22 renal carcinoma patients who exhibited no evidence of CKD (eGFR < 60 mL/min/1.73 m2 or detected dipstick albuminuria). These nephrectomy control group samples were also stored in RNAlater immediately after surgical excision via the same protocol used for the biopsy core tissues. The frozen cortical tissues from the nephrectomy controls were initially cut into the size of a biopsy core, and all collected kidney tissues were manually microdissected under a stereomicroscope (Olympus, Japan) to remove glomeruli. After thorough removal of glomeruli, total RNA was extracted from the remaining tubulointerstitial tissues using an RNeasy Micro Kit (Qiagen, Hamburg, Germany). |

#### Supplementary Table 5: TLR genes pan-disease transcriptomic changes.

| Gene | Group | Log2FC | P value | p.adj |
|---|---|---|---|---|
| | Exp 3-4hr_whole_kidney | 1.52261046 | 0.0040402 | 0.04748356 |
| | Exp 7-8hr_whole_kidney | 0.92826276 | 0.10146881 | 0.25379422 |
| | DN - Adv_whole_kidney | 0.70971932 | 5.7714E-06 | 3.4107E-05 |
| | IgAN - Glom | 1.45069531 | 0.00150326 | 0.00835348 |
| | IgAN - TI | 0.47496259 | 0.00357766 | 0.02562322 |
| | FSGS - TI | 1.4841272 | 0.00011263 | 0.00164522 |
| | LN - TI | 1.7889272 | 3.1739E-06 | 0.00024328 |
| | MN - TI | 1.05635361 | 0.00188692 | 0.04562147 |
| **TLR2** | MCD - TI | 0.73267844 | 0.00313222 | 0.05257494 |
| | Exp 3-4hr_whole_kidney | 1.55361686 | 0.00030413 | 0.00892143 |
| | Exp 7-8hr_whole_kidney | 1.37276985 | 0.00282557 | 0.02214834 |
| | DN - Adv_whole_kidney | 0.5102827 | 4.164E-06 | 2.5541E-05 |
| | IgAN - Glom | 0.38463107 | 0.1217056 | 0.22863175 |
| | IgAN - TI | 0.33488587 | 0.02415035 | 0.09379977 |
| | FSGS - TI | 1.28135946 | 0.00026808 | 0.00310751 |
| | LN - TI | 1.18092974 | 0.00078378 | 0.01667149 |
| | MN - TI | 0.73107802 | 0.01866889 | 0.15437059 |
| **TLR4** | MCD - TI | 0.04958863 | 0.82681911 | 0.9362462 |
| | Exp 3-4hr_whole_kidney | 2.2128265 | 0.00013063 | 0.00507247 |
| | Exp 7-8hr_whole_kidney | 2.22212034 | 0.00029904 | 0.00452622 |
| | DN - Adv_whole_kidney | 1.95725595 | 2.8049E-11 | 6.4609E-10 |
| | IgAN - Glom | 1.3969705 | 0.06595984 | 0.14435816 |
| | IgAN - TI | 1.18512394 | 4.6122E-05 | 0.00116819 |
| | FSGS - TI | 1.91953061 | 0.00444896 | 0.02582251 |
| | LN - TI | 3.60933858 | 7.1042E-08 | 1.1674E-05 |
| | MN - TI | 2.33698688 | 8.58E-05 | 0.00763459 |
| **TLR8** | MCD - TI | 0.54942486 | 0.21337235 | 0.53574642 |

#### Supplementary Table 6: Mitochondrial function related genes pan-disease transcriptomic changes.

| gene | dataset_label | log2FoldChange | pvalue | padj |
|---|---|---|---|---|
| | Exp 3-4hr_whole_kidney | 5.79547687 | NA | NA |
| | Exp 7-8hr_whole_kidney | -0.72100348 | NA | NA |
| | DN - Adv_whole_kidney | -3.53457425 | 1.1542E-11 | 2.9367E-10 |
| | IgAN - Glom | -0.36404786 | 0.04703768 | 0.11252718 |
| | IgAN - TI | -0.66730707 | 2.2953E-06 | 0.00011433 |
| | FSGS - TI | -1.11476454 | 0.00088466 | 0.00758282 |
| | LN - TI | -1.55792668 | 3.3745E-06 | 0.00025498 |
| | MN - TI | -1.42499636 | 1.4918E-06 | 0.00054937 |
| **MT-ATP6** | MCD - TI | -1.23240332 | 1.0822E-08 | 5.6506E-06 |
| | Exp 3-4hr_whole_kidney | -1.27095471 | NA | NA |
| | Exp 7-8hr_whole_kidney | -23.2588375 | NA | NA |
| | DN - Adv_whole_kidney | -2.56370763 | 2.5934E-15 | 1.71E-13 |
| | IgAN - Glom | -0.49589736 | 0.01168477 | 0.04007014 |
| | IgAN - TI | -0.74083506 | 1.7528E-07 | 1.4167E-05 |
| | FSGS - TI | -1.41899825 | 2.507E-05 | 0.00053965 |
| | LN - TI | -1.56455023 | 3.3772E-06 | 0.00025498 |
| | MN - TI | -1.54415353 | 2.0753E-07 | 0.00010137 |
| **MT-CO2** | MCD - TI | -1.1850616 | 4.3945E-08 | 1.5878E-05 |
| | Exp 3-4hr_whole_kidney | -0.7240668 | NA | NA |
| | Exp 7-8hr_whole_kidney | -8.06194458 | NA | NA |
| | DN - Adv_whole_kidney | -1.83910772 | 4.1581E-12 | 1.1979E-10 |
| | IgAN - Glom | -0.77925012 | 0.00169032 | 0.00914005 |
| | IgAN - TI | -0.73080029 | 3.6332E-07 | 2.5193E-05 |
| | FSGS - TI | -1.63518638 | 1.6333E-06 | 7.8599E-05 |
| | LN - TI | -1.74857065 | 2.9499E-07 | 3.8428E-05 |
| | MN - TI | -1.56719916 | 1.9646E-07 | 9.7968E-05 |
| **MT-CO3** | MCD - TI | -1.18062946 | 7.2812E-08 | 2.4817E-05 |
| | Exp 3-4hr_whole_kidney | 0.32044828 | 0.20816861 | 0.43351251 |
| | Exp 7-8hr_whole_kidney | 0.00196592 | 0.99423711 | 0.99741092 |
| | DN - Adv_whole_kidney | -2.58312374 | 6.9028E-15 | 4.0715E-13 |
| | IgAN - Glom | -0.5331663 | 0.02016573 | 0.06025044 |
| | IgAN - TI | -0.61366274 | 2.6771E-05 | 0.00077092 |
| | FSGS - TI | -1.30693161 | 0.00016538 | 0.00217613 |
| | LN - TI | -1.57038686 | 6.0092E-06 | 0.00041503 |
| | MN - TI | -1.43036284 | 3.0442E-06 | 0.00094888 |
| **MT-CYB** | MCD - TI | -1.36273865 | 1.0019E-09 | 8.7549E-07 |
| | Exp 3-4hr_whole_kidney | -0.31911864 | 0.84617115 | NA |
| | Exp 7-8hr_whole_kidney | -0.49967138 | 0.77908387 | NA |
| | DN - Adv_whole_kidney | -2.95051436 | 4.8985E-23 | 2.4537E-20 |
| | IgAN - Glom | -0.23381592 | 0.40643036 | 0.54545504 |
| | IgAN - TI | -0.44273147 | 0.00182879 | 0.01601426 |
| | FSGS - TI | -1.10317728 | 0.0010722 | 0.00877644 |
| | LN - TI | -1.47724004 | 1.1873E-05 | 0.0006894 |
| | MN - TI | -1.44359345 | 1.2577E-06 | 0.00047784 |
| **MT-ND3** | MCD - TI | -1.02062533 | 2.515E-06 | 0.00037633 |
| | Exp 3-4hr_whole_kidney | 0.10134013 | 0.65915465 | 0.80932342 |
| | Exp 7-8hr_whole_kidney | -0.16327775 | 0.50620729 | 0.69051659 |
| | DN - Adv_whole_kidney | -3.11591813 | 1.4482E-10 | 2.7821E-09 |
| | IgAN - Glom | -0.36943241 | 0.04738707 | 0.11317875 |
| | IgAN - TI | -0.60567501 | 8.5586E-06 | 0.00032588 |
| | FSGS - TI | -1.11787065 | 0.00054064 | 0.00531756 |
| | LN - TI | -1.34822637 | 3.01E-05 | 0.00139369 |
| | MN - TI | -1.31038953 | 4.3887E-06 | 0.00118032 |
| **MT-ND4** | MCD - TI | -1.03550063 | 6.1996E-07 | 0.00013125 |

**Supplementary Table 7: Consistently Upregulated TLR related pathways across podocyte ablated mice and human glomerulus dysfunction related disease cohorts via GSEA analysis**

| Description | dataset_label | NES | pvalue | qvalue | setSize | sig_label |
|---|---|---|---|---|---|---|
| GOBP_TOLL_LIKE_RECEPTOR_SIGNALING_PATHWAY | Exp_3_4_hr_Whole_kidney | 1.89399306 | 9.4447E-06 | 8.2751E-05 | 79 | *** |
| | DN_Whole_kidney | 1.62597531 | 0.00054188 | 0.00736559 | 84 | ** |
| | LN_TI | 2.5158784 | 1E-10 | 3.4331E-09 | 82 | *** |
| | MN_TI | 1.95300165 | 1.8534E-05 | 0.00048653 | 82 | *** |
| | MCD_TI | 1.63419481 | 0.00136447 | 0.02675577 | 82 | * |
| GOBP_MYD88_DEPENDENT_TOLL_LIKE_RECEPTOR_SIGNALING_PATHWAY | Exp_3_4_hr_Whole_kidney | 1.63194151 | 0.01000132 | 0.02397471 | 21 | * |
| | IgAN_Glom. | 2.42540129 | 2.372E-05 | 0.00384759 | 20 | ** |
| | FSGS_TI | 1.91479577 | 0.00076297 | 0.00523396 | 31 | ** |
| | LN_TI | 2.15417867 | 1.2829E-05 | 0.00015707 | 21 | *** |
| | MN_TI | 1.98329539 | 0.00037142 | 0.00612998 | 21 | ** |
| | MCD_TI | 1.85830403 | 0.00216922 | 0.03717008 | 21 | * |
| GOBP_ENDOLYSOSOMAL_TOLL_LIKE_RECEPTOR_SIGNALING_PATHWAY | Exp_3_4_hr_Whole_kidney | 1.70938545 | 0.00088018 | 0.00357948 | 55 | ** |
| | IgAN_TI | 1.92716163 | 8.4794E-05 | 0.00369545 | 52 | ** |
| | LN_TI | 2.33078072 | 6.2818E-09 | 1.5801E-07 | 52 | *** |
| | MN_TI | 2.21896756 | 4.0521E-07 | 1.6963E-05 | 52 | *** |
| | MCD_TI | 1.92978481 | 0.00020009 | 0.00677384 | 52 | ** |

**Supplementary Table 8: Consistently downregulated mitochondria function-related pathways across podocyte-ablated mice and human glomerulus dysfunction-related disease cohorts via GSEA analysis.**

| Description | dataset_label | NES | pvalue | qvalue | setSize | sig_label |
|---|---|---|---|---|---|---|
| **HP_MITOCHONDRIAL_INHERITANCE** | Exp_3_4_hr_Whole_kidney | -1.94944 | 0.001271 | 0.004788 | 15 | ** |
| | Exp_7_8_hr_Whole_kidney | -2.1531 | 9.39E-08 | 6.39E-05 | 15 | *** |
| | DN_Whole_kidney | -2.9082 | 1E-10 | 1.13E-08 | 28 | *** |
| | FSGS_TI | -1.74991 | 0.003495 | 0.018745 | 28 | * |
| | LN_TI | -2.00112 | 0.000161 | 0.001476 | 28 | ** |
| | MN_TI | -2.21944 | 1.6E-06 | 5.62E-05 | 28 | *** |
| | MCD_TI | -2.03152 | 1.71E-05 | 0.001063 | 28 | ** |
| **HP_MITOCHONDRIAL_RESPIRATORY_CHAIN_DEFECTS** | Exp_7_8_hr_Whole_kidney | -1.97043 | 1.63E-05 | 0.003699 | 18 | ** |
| | DN_Whole_kidney | -2.58605 | 6.35E-10 | 6.13E-08 | 19 | *** |
| | LN_TI | -1.87867 | 0.001564 | 0.010313 | 19 | * |
| | MN_TI | -2.04007 | 0.000155 | 0.002974 | 19 | ** |
| | MCD_TI | -1.87621 | 0.000477 | 0.013014 | 19 | * |
| **HP_ABNORMAL_MITOCHONDRIA_IN_MUSCLE_TISSUE** | Exp_3_4_hr_Whole_kidney | -1.995 | 0.00035 | 0.001687 | 39 | ** |
| | Exp_7_8_hr_Whole_kidney | -2.48272 | 1E-10 | 2.31E-07 | 39 | *** |
| | DN_Whole_kidney | -2.38431 | 1.27E-07 | 6.2E-06 | 47 | *** |
| | IgAN_TI | -1.70769 | 0.002573 | 0.042025 | 47 | * |
| | FSGS_TI | -2.00159 | 0.000235 | 0.001929 | 47 | ** |
| | LN_TI | -1.84827 | 0.000403 | 0.00325 | 47 | ** |
| | MN_TI | -2.17145 | 1.01E-05 | 0.000291 | 47 | *** |
| | MCD_TI | -1.92553 | 6.53E-05 | 0.002897 | 47 | ** |
| **HP_MITOCHONDRIAL_MYOPATHY** | Exp_3_4_hr_Whole_kidney | -1.68786 | 0.003783 | 0.011427 | 54 | * |
| | Exp_7_8_hr_Whole_kidney | -2.4474 | 1E-10 | 2.31E-07 | 54 | *** |
| | DN_Whole_kidney | -2.7838 | 1E-10 | 1.13E-08 | 63 | *** |
| | IgAN_TI | -1.79041 | 0.000453 | 0.012675 | 63 | * |
| | FSGS_TI | -1.82334 | 0.000589 | 0.004231 | 63 | ** |
| | LN_TI | -1.73544 | 0.001689 | 0.011008 | 63 | * |
| | MN_TI | -2.2424 | 1.12E-07 | 5.54E-06 | 63 | *** |
| | MCD_TI | -1.81362 | 0.000244 | 0.007985 | 63 | ** |
| **GOBP_MITOCHONDRIAL_ELECTRON_TRANSPORT_CYTOCHROME_C_TO_OXYGEN** | Exp_7_8_hr_Whole_kidney | -2.12516 | 1.74E-05 | 0.003807 | 19 | ** |
| | DN_Whole_kidney | -2.04783 | 0.00057 | 0.007669 | 20 | ** |
| | IgAN_TI | -2.1283 | 1.14E-05 | 0.000707 | 20 | *** |
| | FSGS_TI | -2.20369 | 4.02E-05 | 0.000406 | 20 | *** |
| | LN_TI | -1.77719 | 0.007283 | 0.036713 | 20 | * |
| | MN_TI | -1.95983 | 0.000335 | 0.005587 | 20 | ** |
| **HP_MUSCLE_ABNORMALITY_RELATED_TO_MITOCHONDRIAL_DYSFUNCTION** | Exp_7_8_hr_Whole_kidney | -2.30173 | 5.7E-09 | 7.33E-06 | 87 | *** |
| | DN_Whole_kidney | -2.61817 | 1E-10 | 1.13E-08 | 100 | *** |
| | IgAN_TI | -1.80691 | 0.000174 | 0.006318 | 100 | ** |
| | FSGS_TI | -1.71648 | 0.001104 | 0.007147 | 100 | ** |
| | LN_TI | -1.63066 | 0.002655 | 0.015916 | 100 | * |
| | MN_TI | -2.12447 | 7.76E-07 | 2.93E-05 | 100 | *** |
| | MCD_TI | -1.6447 | 0.001195 | 0.024789 | 100 | * |

**Supplementary Table 9: Consistently Upregulated Apoptosis pathways across podocyte-ablated mice and human glomerulus dysfunction-related disease cohorts via GSEA analysis**

| Description | dataset_label | NES | pvalue | qvalue | setSize | sig_label |
|---|---|---|---|---|---|---|
| GOBP_REGULATION_OF_EXTRINSIC_APOPTOTIC_SIGNALING_PATHWAY | Exp_3_4_hr_Whole_kidney | 1.97407 | 4.71E-09 | 1.13E-07 | 157 | *** |
| | IgAN_TI | 1.524647 | 0.00261 | 0.042311 | 167 | * |
| | LN_TI | 1.826985 | 7.53E-06 | 9.85E-05 | 167 | *** |
| | MN_TI | 1.49422 | 0.001601 | 0.019791 | 167 | * |
| | MCD_TI | 1.642373 | 0.000126 | 0.00471 | 167 | ** |
| GOBP_EXTRINSIC_APOPTOTIC_SIGNALING_PATHWAY | Exp_3_4_hr_Whole_kidney | 2.071804 | 1E-10 | 3.81E-09 | 225 | *** |
| | LN_TI | 1.931739 | 1.18E-08 | 2.83E-07 | 239 | *** |
| | MN_TI | 1.431781 | 0.002525 | 0.028912 | 239 | * |
| | MCD_TI | 1.565392 | 9.51E-05 | 0.003852 | 239 | ** |
| GOBP_APOPTOTIC_CELL_CLEARANCE | Exp_3_4_hr_Whole_kidney | 1.672703 | 0.002102 | 0.007155 | 37 | ** |
| | LN_TI | 2.300509 | 4.68E-08 | 1.01E-06 | 44 | *** |
| | MN_TI | 2.087208 | 7.85E-06 | 0.000229 | 44 | *** |
| | MCD_TI | 1.792683 | 0.001601 | 0.03031 | 44 | * |

**Supplementary Table 10: Primary antibodies used for the immunohistochemistry studies.**

| PRIMARY ANTIBODY | SPECIES | COMPANY | CATALOG | DILUTION | SECONDARY ANTIBODY | COLOR DEVELOPMENT |
|---|---|---|---|---|---|---|
| CD59 | Rabbit | Sigma | HPA026494 | 1:200 | OMap anti-Rabbit HRP | Discovery Rhodamine |
| Synaptopodin | Rabbit | ProteinTech | 21064-1-AP | 1:400 | OMap anti-Rabbit HRP | Discovery Cy5 |
| Dapi | | Invitrogen | D1306 | | | |
